# CCAR1 promotes DNA repair via an unanticipated role in alternative splicing

**DOI:** 10.1101/2023.09.18.557780

**Authors:** Mehmet E. Karasu, Brian J. Joseph, Aleksei Mironov, Markus S. Schroeder, Ana Gvozdenovic, Mihaela Zavolan, Jacob E. Corn

**Affiliations:** Department of Biology, ETH Zurich, Zurich 8053, Switzerland; Department of Systems Biology, Department of Biochemistry and Molecular Biophysics, Center for Motor Neuron Biology and Disease, Columbia University, New York NY 10032, USA; Computational and Systems Biology, Biozentrum, University of Basel, 4056, Basel, Switzerland

## Abstract

DNA repair is directly performed by hundreds of core factors and indirectly regulated by thousands of others. We massively expanded a CRISPR inhibition and Cas9-editing screening system to discover factors indirectly modulating homology directed repair (HDR) in the context of ∼18’000 individual gene knockdowns. We focused on *CCAR1,* a poorly understood gene that we found reduced both HDR and interstrand crosslink repair, phenocopying the loss of the Fanconi Anemia pathway. *CCAR1* loss abrogated FANCA protein without substantial reduction in the level of its mRNA or that of other FA genes. We instead found that CCAR1 prevents inclusion of a poison exon in *FANCA*. Transcriptomic analysis revealed that the CCAR1 splicing modulatory activity is not limited to FANCA, and it instead regulates widespread changes in alternative splicing that would damage coding sequences in mouse and human cells. CCAR1 therefore has an unanticipated function as a splicing fidelity factor.

## Introduction

Eukaryotic DNA repair is a complex process directly involving approximately 500 known proteins that work in inter-dependent and overlapping pathways^1^. Double-stranded DNA break (DSB) repair involves multiple possible routes that can either be error-prone or precise. Failure to repair a DSB is particularly dangerous as it would leave the resulting DNA without either a centromere or telomeres, compromising a great deal of genetic information.

Eukaryotes have developed two major paths to DSB repair: untemplated non-homologous end joining (NHEJ) and templated homology directed repair (HDR), each encompassing multiple sub-pathways with their own set of facilitating factors^2–4^. DSB repair has attracted a great deal of interest in recent years, not only for its fundamental importance in genome biology, but also because it is harnessed during CRISPR-Cas genome editing^5,6^

While hundreds of proteins are directly involved in DNA repair, thousands of other factors that regulate these processes indirectly. For example, HDR is most active during the S/G2 phases of the cell cycle, in part through post-translational modification of DNA repair proteins by cell cycle regulated kinases ^7,8^. Cell cycle regulation thereby indirectly alters a cell’s propensity to perform HDR ^9^. Epigenetic modifications can also indirectly alter repair propensities by regulating the availability of a given segment of DNA to various repair proteins ^10^. Exploring how broader cell biology connects to DNA repair therefore helps understand emergent genetic properties and fuels advances in genome editing.

We previously performed limited screens (∼2’000 genes) that coupled CRISPR transcriptional inhibition (CRISPRi) to a precise Cas9-induced DSB, using flow cytometry to identify genes involved in human HDR ^11,12^. Our prior work found a crucial role for the Fanconi Anemia (FA) pathway in CRISPR-Cas HDR and identified factors whose removal promotes higher HDR.

Here, we massively expanded our search to genome-wide (∼18’000 genes) identification of unanticipated moonlighting factors not typically associated with DNA repair. Our approach re-identified repair factors known to be critical for DSB repair, validating the scaled-up approach. We also found many new genes with previously unknown connections to DNA repair. We chose to focus our mechanistic investigation on *CCAR1*, a poorly understood gene with a known role in the regulation of transcript abundance. We found that *CCAR1* expression is required for both HDR and interstrand crosslink (ICL) repair by ensuring the protein expression of the FA-pathway member FANCA. Surprisingly, knockdown of *CCAR1* reduces FANCA protein levels without much affecting its transcript levels. Transcriptomic studies revealed that CCAR1 represses the inclusion of a FANCA poison exon, as well as regulating more than 300 damaging exon inclusion and exclusion events in human and mouse cells. Our data thus indicate that CCAR1 has a broad and conserved, though so far unanticipated, role as a splicing fidelity factor.

## Results

### CCAR1 regulates HDR and interstrand crosslink repair

To identify human genes involved in CRISPR-Cas-mediated HDR, we conducted a genome-wide screen in K562 cells using our previously described CRISPRi-coupled CRISPR nuclease system ^11–13^, as illustrated in **Figure S1A** (see Methods). Consistent with published studies, we identified genes associated with the FA pathway as high-confidence hits that promote HDR (**Figure 1A, Table 1)**. An unbiased gene ontology analysis of all high-confidence hits from the genome-wide screen revealed an enrichment in relevant pathways, such as DSB repair via homologous recombination (**Figure S1B).** These findings indicate that the genome-wide screen identified established participants in DNA damage response (DDR) and gave us confidence in its capability to uncover novel participants in DDR.

**Figure 1.**
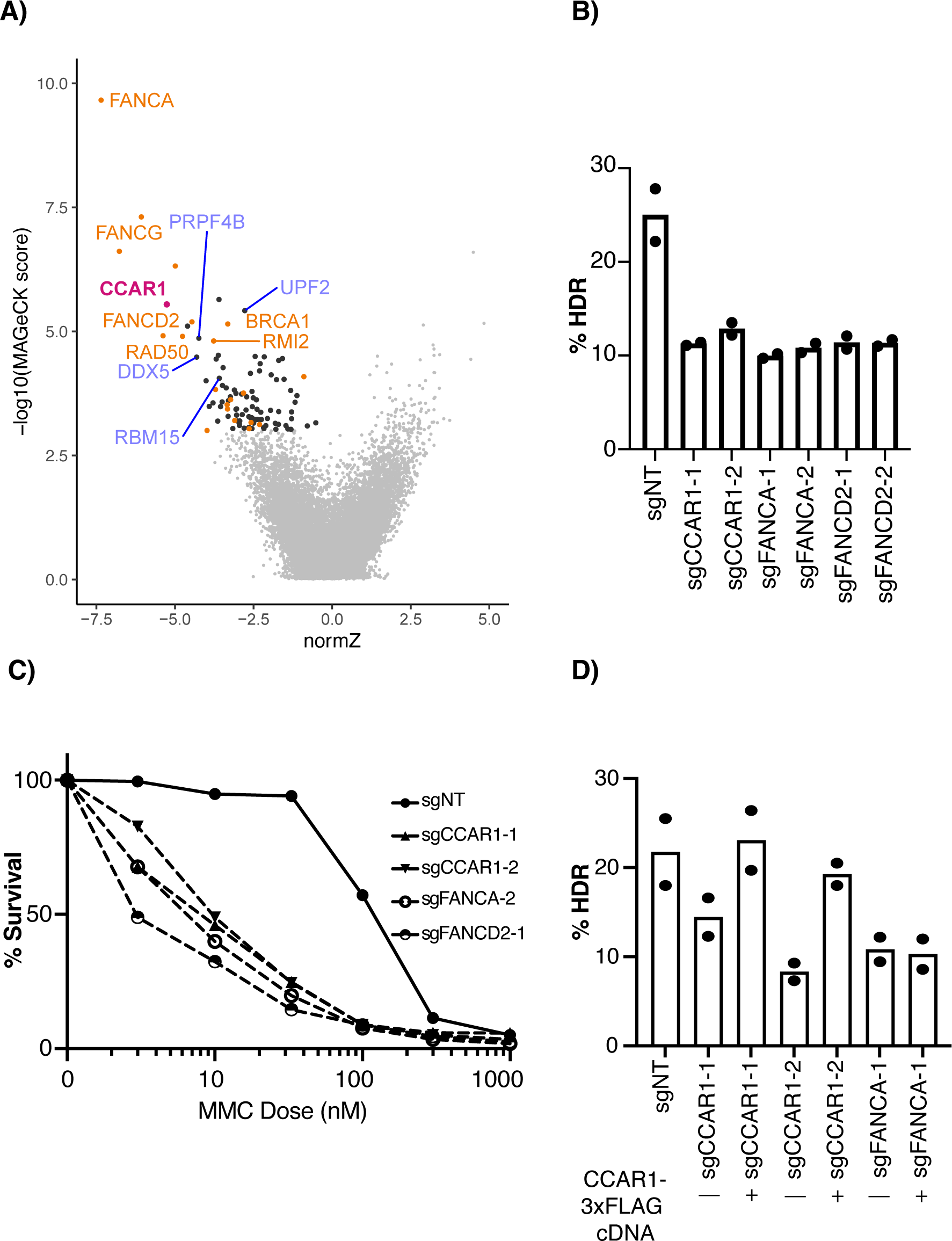
Knockdown of *CCAR1* phenocopies FA pathway deficiency. **A)** Analysis of a genome-wide coupled CRISPRi and Cas9 HDR screen. Gene-level abundance was compared in GFP-positive (HDR proficient) vs. unsorted background population using MaGECK and DrugZ (see Table 1). Genes highlighted in orange are selected HDR repair factors identified in previous screens, while genes highlighted in blue are associated with splice regulation. **B)** CRISPRi knockdown of *CCAR1* reduces CRISPR-Cas9-mediated HDR in K562 cells. Clonal CRISPRi stable cell lines were individually transduced with two individual lentiviral sgRNAs targeting *CCAR1*, *FANCA*, and *FANCD2*. A BFP-to-GFP HDR assay was used to measure HDR efficiency. Each dot represents an individual biological replicate, and the bars represent the mean. **C)** *CCAR1* knockdown cells display increased MMC sensitivity. The black constant line indicates control cells transduced with non-targeting (NT) guide RNA, while the dashed lines with different shapes represent *CCAR1*, *FANCA*, or *FANCD2* knockdown cells, respectively. **D)** Addition of *CCAR1* cDNA rescues HDR deficiency in *CCAR1* knockdown cells. K562 CRISPRi cell lines were stably transduced or not with CCAR1-3xFLAG cDNA, as indicated. The BFP-to-GFP assay was used to measure HDR efficiency in each cell line. Each dot represents an individual biological replicate and the bars represent the mean.

We observed a striking reduction in CRISPR-Cas-mediated HDR upon depletion of Cell Cycle and Apoptosis Regulator 1 (*CCAR1*) (**Figure 1A)**. *CCAR1* is a poorly understood gene that encodes a 1150-amino-acid-long protein, originally identified in a screen for genes responsible for retinoic acid-mediated apoptosis in human breast cancer cells ^14^. CCAR1 has been recognized for its involvement in the Mediator complex, thereby regulating the transcription of genes involved in pathways ranging from Beta-catenin signaling to adipocyte differentiation ^15–18^. A recent CRISPR screen for genes involved in resistance to diverse DNA-damaging agents found that *CCAR1* knockout leads to interstrand crosslink hypersensitivity ^16^. However, mechanistic understanding remained elusive and a role of CCAR1 in HDR was not described.

We cloned individual CRISPRi guide RNAs targeting *CCAR1* into a lentiviral expression construct, as well as guides targeting FA pathway members such as *FANCA* and *FANCD2*. Knocking down *CCAR1* led to a reduction in HDR levels in a BFP-to-GFP assay similar to knockdown of *FANCA* and *FANCD2* (**Figure 1B**). *CCAR1* knockdown also resulted in increased sensitivity to the DNA crosslinker mitomycin C (MMC) (**Figure 1C**). These results were consistent across multiple guide RNAs for both *CCAR1* and the FA genes (**Figure S2A, B, C**). We conducted rescue experiments by introducing *CCAR1* into cells with either CCAR1 or FANCA CRISPRi-mediated depletion. We found that HDR efficiency was rescued by introducing *CCAR1* cDNA to *CCAR1* knockdown cells, but knockdown of *FANCA* could not be rescued by introducing *CCAR1* (**Figure 1D**), suggesting that FANCA might be genetically downstream of CCAR1. These data strongly indicate that *CCAR1* promotes HDR and reveal an unexpected yet important connection between CCAR1 and multiple functions of the FA pathway.

### CCAR1 regulates the abundance of the FANCA protein but not mRNA

To identify the specific dysfunction that *CCAR1* loss causes within the FA pathway, we tested the abundance and molecular activity of various FA pathway members ^17,18^. Control cells targeted by a non-targeting guide RNA were competent for mono-ubiquitination of FANCD2 upon MMC treatment (**Figure 2A**), and as expected, the knockdown of *FANCA* abolished this activity. Knockdown of *CCAR1* was also sufficient to abolish MMC-induced FANCD2 monoubiquitination.

**Figure 2.**
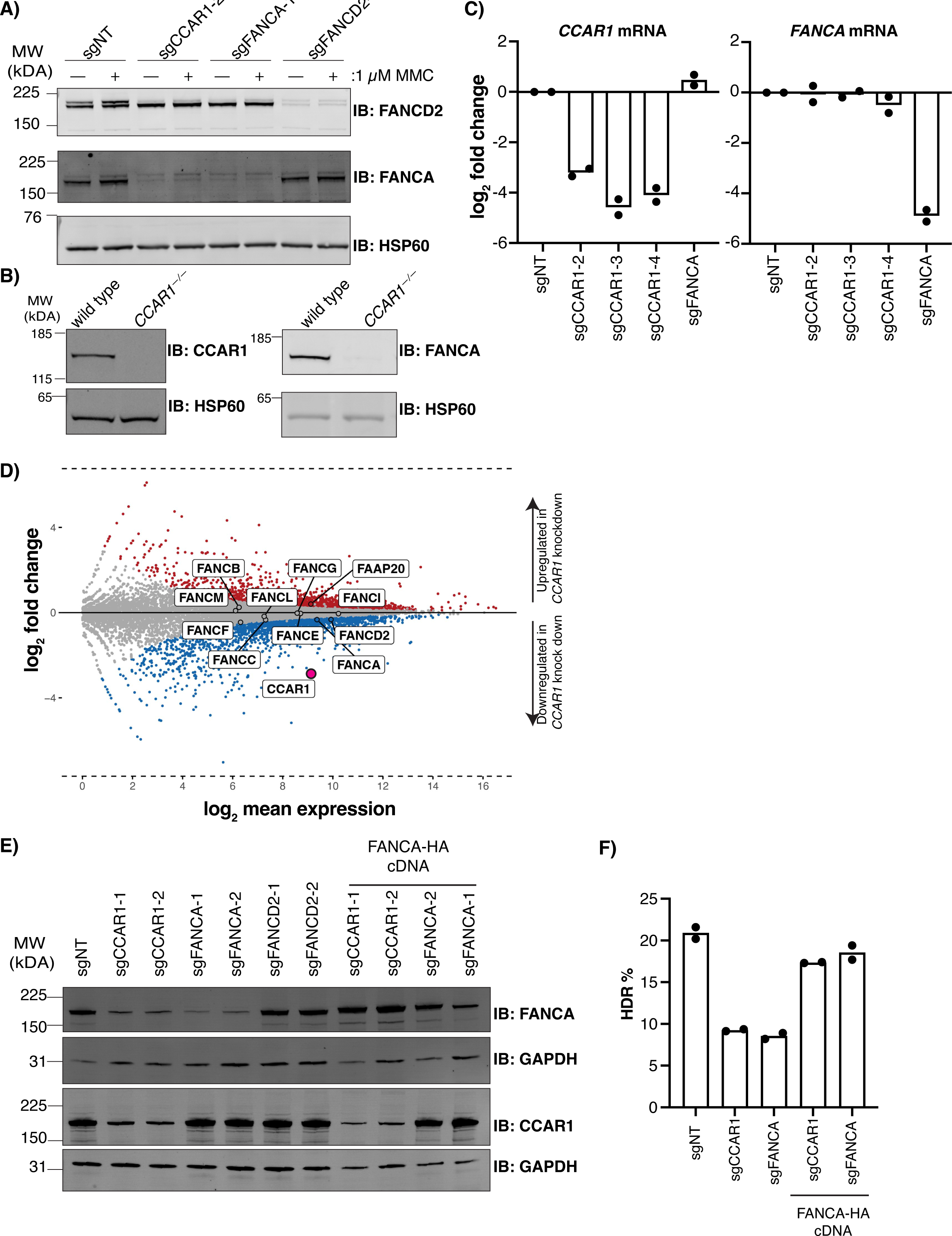
CCAR1 regulates FANCA protein stability without affecting its transcript level. **A)** The FANCA protein is depleted in the absence of *CCAR1*. K562 cells with various stable CRISPRi knockdowns were exposed to 1 μM MMC for 24 h. Protein extracts were analyzed by Western blot using indicated antibodies (anti-FANCA, anti-FANCD2, anti-HSP60). FANCD2 was detected as a doublet, and the slowly migrating FANCD2 band represents FANCD2-Ub. **B)** FANCA is also lost in CCAR1 knockout cells. Western blots comparing *CCAR1^-/-^* K562 cells to wild-type cells probed with anti-CCAR1 and anti-FANCA antibodies. HSP60 was used as a loading control. **C)** *FANCA* mRNA levels are not affected by *CCAR1* knockdown in K562 cells. RT-qPCR analysis of RNAs extracted from the indicated stable CRISPRi cell lines. The plotted values represent the log2 fold difference normalized to non-targeting control (sgNT) sample. Two independent biological replicates were conducted and are represented as dots, with bars indicating the means for each cell line. **D)** RNA-seq reveals minimal transcriptional changes in FA pathway genes in *CCAR1* CRISPRi vs. wild-type K562 cells. The orange magenta dot indicates *CCAR1* and the core FA genes are highlighted. **E, F)** Ectopic expression of *FANCA* cDNA rescues HDR in CCAR1 stable CRISPRi K562 cells. Protein extracts were analyzed by Western blot using the indicated antibodies (anti-FANCA, ant-CCAR1, anti-GAPDH1). Indicated cells carrying stably expressed sgRNAs and stably overexpressed FANCA cDNA were subjected to the BFP-to-GFP HDR assay. Each dot represents an individual biological replicate and bars represent the mean.

Knockdown of *FANCA* and *FANCD2* abolished expression of each respective protein (**Figure 2A**). Unexpectedly, *CCAR1* knockdown reduced the levels of FANCA protein without affecting FANCD2 protein levels (**Figure 2A)**. We verified that depleting *CCAR1* leads to loss of FANCA protein by generating isogenic CRISPR-Cas9-mediated *CCAR1* knockout clones in K562 cells (**Figure 2B**). Western blots revealed that *CCAR1^-/-^* cells also lacked FANCA protein expression (**Figure 2B**), confirming that CCAR1 regulates FANCA protein production and/or stability.

Prior studies suggested that CCAR1 acts as a transcriptional co-factor ^15,19,20^, so we first investigated if *CCAR1* regulates *FANCA* transcription. However, we found that the level of *FANCA* mRNA was largely unaffected in *CCAR1* knockdown K562 cells (**Figure 2C**). This was also true in retinal pigment epithelial (RPE1) cells, where depletion of *CCAR1* also decreased HDR efficiency and abolished FANCA protein expression without grossly reducing its mRNA level (**Figure S2D-F**).

We next investigated potential post-transcriptional regulation of *FANCA* abundance. The stability of FANCA relies on interactions with other FA core complex members, such as FAAP20 and FANCG ^21,22^. We therefore asked if *CCAR1* regulates the transcription of other FA genes, thereby indirectly affecting FANCA protein levels. We performed RNA-seq in wildtype and *CCAR1*-knockdown K562 cells. Consistent with CCAR1’s known roles in transcription, *CCAR1* knockdown impacted the expression of numerous genes (2’211 upregulated, 2’064 downregulated, FDR < 0.05) (**Figure 2D, S3**). However, *CCAR1* status did not greatly affect *FANCA* mRNA levels, nor did it induce substantial changes in the transcription of other FA pathway genes (**Figure 2D, Table 2**).

While performing cDNA complementation to test the role of CCAR1 in FA activity and transcription, we found that re-expression of *FANCA* cDNA in *CCAR1* knockdown cells could rescue both FANCA expression and HDR (**Figure 2E, F**). This was surprising, because our transcriptional data indicated that the *FANCA* mRNA was still present in *CCAR1* knockdown cells (**Figure 2C, D**). Therefore, in *CCAR1*-depleted cells the ectopic *FANCA* cDNA is competent for FANCA protein and RNA expression, while the endogenous FANCA mRNA is somehow compromised.

### CCAR1 represses inclusion of a poison exon in *FANCA* mRNA

Closely examining our RNA-seq data, we found that the *CCAR1* knockdown induced a novel splicing event within the *FANCA* mRNA (**Figure 3A, S4A**). Specifically, we identified a cassette exon (CE) within FANCA, positioned between the conventional exons 14 and 15 (chr16:89,785,124-89,785,159, hg38). While this 36 bp CE would not alter the reading frame, it contains an in-frame stop codon at the 3’ end and is therefore a poison exon (PE). The PE is included in most sequenced transcripts and would lead to the premature termination of FANCA translation, consistent with the loss of FANCA protein expression (**Figure 3A**). Using RT-PCR, we further validated that this *FANCA* PE is efficiently spliced into the mRNA in two independent CCAR1 knockout clones but not the isogenic wildtype, and this splicing is reversed by cDNA re-expression of *CCAR1* in the knockout background. (**Figure 3B, Figure S4B)**.

**Figure 3.**
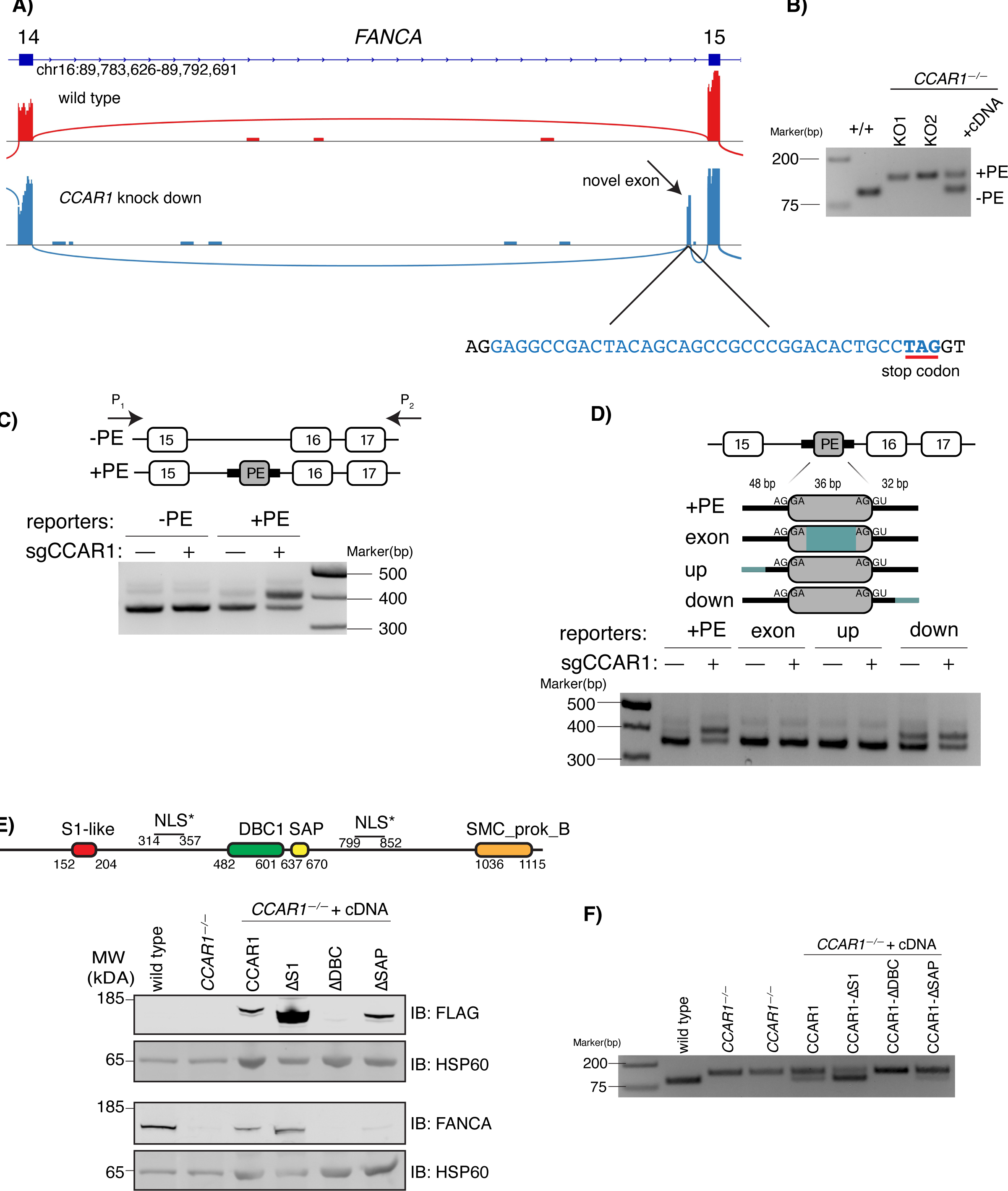
A CCAR-1regulated novel poison exon leads to non-functional *FANCA* mRNA. **A)** CCAR1 knockdown leads to splicing of a novel cassette exon into the *FANCA* mRNA. Sashimi plots indicate the mapped RNA-seq reads from wild type (red) and *CCAR1* stable CRISPRi (blue) tracks. A novel exon inclusion is highlighted by the arrow. Underneath the sashimi plots, the novel exon sequence is shown in light blue and the stop codon (bold) is underlined. Splicing acceptor and donor sites are shown in black. (Sashimi plots from replicates 2 and 3 are shown in **Figure S4A**). **B)** FANCA mRNA from K562 *CCAR1^-/-^* cells include the poison exon. RT-PCR experiments were performed on the indicated samples: wild type, two different *CCAR1^-/-^* clones, and a *CCAR1^-/-^* clone with stably re-expressed CCAR1 cDNA. The lower band represents the product of wildtype splicing between exon 14 and 15, while the upper band represents the poison exon inclusion. **C)** A mini gene reporter containing the poison exon responds to *CCAR1* presence. Upper, maps of the mini gene constructions. A construct lacking the poison exon (-PE) contains the genomic region of FANCA from exons 15 to 17. A construct containing the poison exon (+PE) was generated by inserting the poison exon surrounding sequences between exon 15 and 16. Bottom, RT-PCR experiments were performed with the primers located in constant parts of the mini-gene reporters. The lower band represents the wildtype spliced product, and the upper band represents the poison exon-included product. **D)** Cis-regulatory elements are required for poison exon inclusion and CCAR1-dependent regulation. Upper, sequences were scrambled in the exonic (30 bp, exon), upstream (14 bp, up), and downstream (17 bp, down) parts of the poison exon. The scrambled sequence information is shown in **Figure S5**. Bottom, RT-PCR experiments were performed as in **panel C** on the indicated samples. The lower band represents the wild-type spliced product, and the upper band represents the poison exon-included product. **E, F)** The S1 RNA binding domain of CCAR1 is dispensable for splicing activity, but the SAP DNA/RNA binding domain is required. Upper, CCAR1 protein domains and predicted NLS sites are indicated (also see **Figure S6A**). Bottom, samples were analyzed by Western blot with the indicated antibodies (anti-FANCA, anti-FLAG, anti-HSP60). **F)** RT-PCR experiments were performed on the same samples as in **panel E** with the addition of another clone of *CCAR1^-/-^* clone. As described previously, the lower band represents the wild-type spliced product while the upper band represents the poison exon-included spliced product.

Going back to our original screening data, we found that several spliceosome associated components regulate CRISPR-Cas HDR such as *PRPF4B*, *RMB15*, *DDX5* (**Figure 1A**). We therefore asked if the FANCA PE is highly specific to *CCAR1* perturbation, or part of a previously unanticipated splicing program that involves other components of the spliceosome or RNA binding proteins. To this end, we re-analyzed ENCODE RNA-seq datasets obtained following the knockdown of various RNA-binding proteins and splicing factors ^23^. We reprocessed the ENCODE datasets (Methods), which tested perturbations for a total of 192 genes in multiple cell lines. We found that inclusion of the *FANCA* PE was induced by several splicing factor perturbations, especially U2AF1 and U2AF2 (**Figure S4C**). These findings underscore that the *FANCA* poison exon inclusion event can be regulated by multiple canonical constituents of the spliceosome, though none of these have as strong effect as *CCAR1* depletion.

To investigate the *cis* nucleic acid elements that regulate CCAR1-mediated *FANCA* splicing, we developed a synthetic splicing assay using a minimal reporter. We transplanted the FANCA PE and an 80 base flanking sequence, normally located between *FANCA* exons 14/15, into a different context between exons 15/16 and electroporated the reporters into K562 cells (**Figure 3C**). Introducing the PE and its surrounding sequence into a new context tests the ability of CCAR1 to regulates its splicing beyond the endogenous context we already observed by RNA-seq. Splicing between exons 15/16 was independent of CCAR1 status in a reporter lacking the PE, and the reporter containing the PE was normally spliced in wild type cells (**Figure 3C**). However, the PE was specifically spliced into the transcript in cells with *CCAR1* knocked down (**Figure 3C**).

We next used the *FANCA* exon 15/16 splicing reporter to identify sequences within and surrounding the *FANCA* PE that dictate its inclusion. We scrambled three sections of the PE and the surrounding sequence: 30 bases within the exon itself, 17 bases upstream, and 14 bases downstream (**Figure 3D and S5**). In all cases, we retained the canonical splicing acceptor and donor sites. Scrambling the PE itself prevented inclusion even in the absence of *CCAR1*, as did scrambling the upstream region (**Figure 3D**). Strikingly, scrambling the downstream region led to constitutive PE inclusion, even in the presence of CCAR1 (**Figure 3D)**. These results indicate that multiple regulatory elements and/or RNA secondary structure modulate the inclusion of this PE, including a downstream element that represses PE inclusion in a CCAR1-dependent manner.

To gain deeper insights into the contribution of specific domains of CCAR1 in regulating *FANCA* PE splicing, we generated multiple deletions of CCAR1 domains ^24^ or fragments and introduced FLAG-tagged constructs into *CCAR1^-/-^* K562 cells. CCAR1 is over 1’000 amino acids in length, and we generated a wide variety of constructs outlined in **Figure S6A.** Here, we focus on the roles of domains with previously describe activities in DNA/RNA metabolism: an S1 RNA binding domain ^25^, a DBC1-like domain that may be involved in transcriptional regulation ^26^, and a SAP domain DNA/RNA binding domain that may be involved in transcription or RNA processing ^27–29^.

We found that the deletion of the S1 RNA binding domain was dispensable for CCAR1-dependent FANCA protein expression (**Figure 3E**). Deletion of the DBC1 domain destabilized CCAR1, preventing the analysis of its potential role in splicing regulation. However, expression of CCAR1 lacking the 33-residue SAP domain did not support FANCA expression. These results were confirmed with an RT-PCR for endogenous *FANCA* splicing, which demonstrated that the SAP domain was required to prevent *FANCA* PE inclusion. (**Figure 3F**). Further deletions also indicated important roles for poorly annotated regions between amino acids 200-314 (N-terminal of the first NLS) and amino acids 950-1000 (N-terminal to an SMC domain) (**Figure S6B**). Although the phosphorylation of amino acid 192 had been deemed significant in certain signalling contexts ^30^, it did not appear essential for this activity in our experiments.

### CCAR1 has a broad splicing regulatory function that is conserved between human and mouse

Having established CCAR1’s role in splicing of the *FANCA* poison exon, we wondered whether it influenced other splicing events. Indeed, MAJIQ analysis of RNA-seq data from K562 cells identified approximately 250 differential exon-intron junction usage events in *CCAR1* knockdown relative to control (confidence cut-off > 0.8, |ΔPSI| > 0.10) ^31^ (**Figure 4A, Table 3**). These alterations primarily stemmed from cassette exon inclusion or exclusion events and included several unannotated novel exons. To determine whether this widespread splicing regulatory activity was evolutionarily conserved, we generated a *Ccar1*-knockout mouse 3T3 fibroblast line (**Figure S7A**) and performed RNA-seq for wild type and *Cccar1^-/-^* cells. Much as in human cells, knockout of *Ccar1* in mouse cells led to approximately 110 differential exon-intron junction usage events, with most of these stemming from cassette exon events (**Figure 4B, Table 4**).

**Figure 4.**
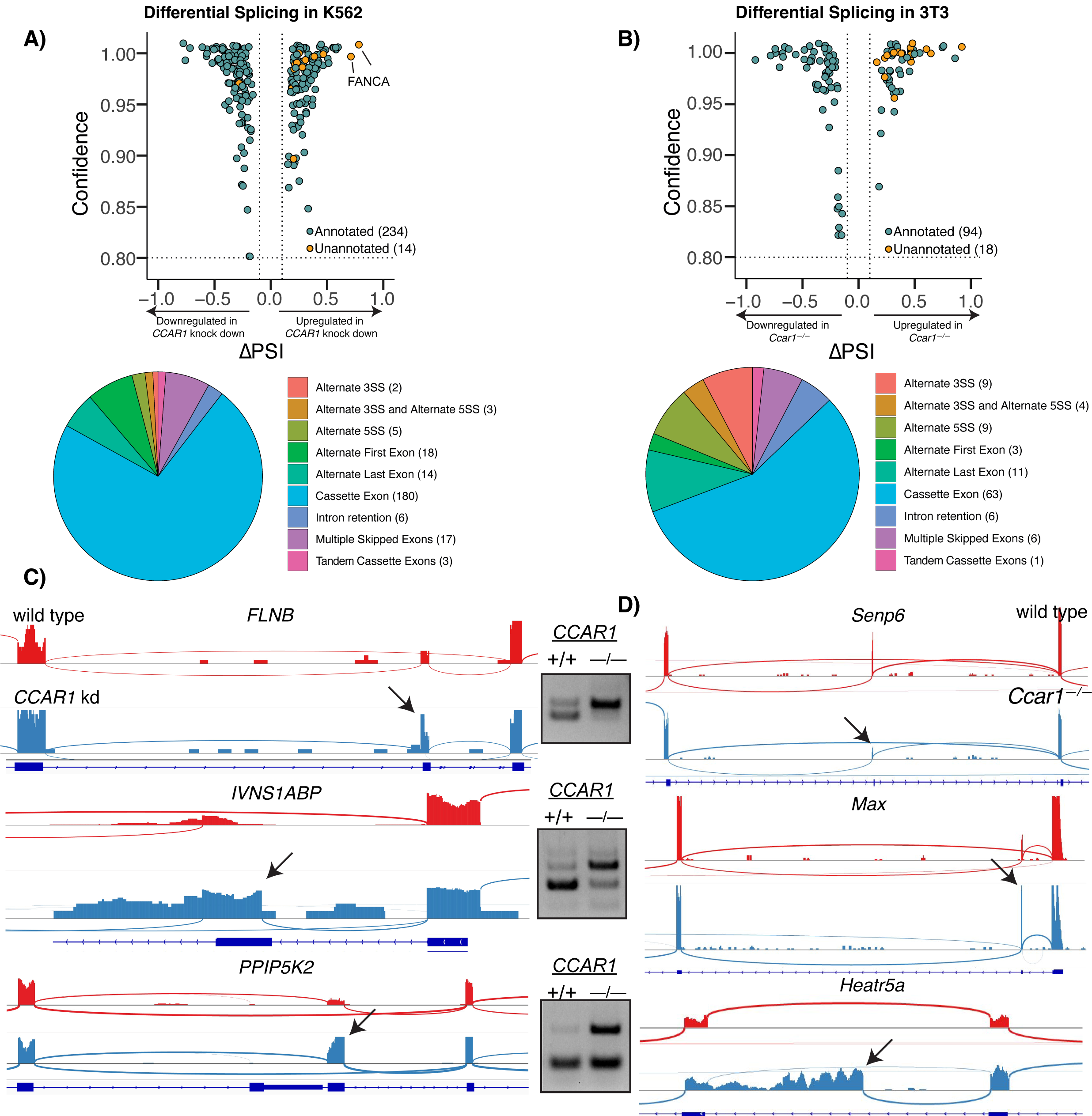
CCAR1 regulates alternative splicing in human and mouse cell lines. **A)** Human *CCAR1* regulates many alternative splicing events in addition to the *FANCA* poison exon. Upper panel: MAJIQ-identified differentially spliced exon-intron junctions in replicate RNA-seq comparing *CCAR1* CRISPRi knockdown and wild-type K562 cells. Orange dots highlight the novel events (unannotated) occurring in the absence of *CCAR1* (full events are in **Table 3**). Bottom panel: Most of the significant CCAR-dependent events relative to cassette exon usage. The number of events in each class is in parentheses. **B)** Mouse *Ccar1* also regulates many alternative splicing events. Analysis was performed for replicate samples *Ccar1^-/-^* and wildtype 3T3 cells, as per **panel A** (full events are in **Table 4**). **C)** Illustration and validation of CCAR-dependent alternative splicing events in human cells. On the left, sashimi plots display the differential exon usage in the noted genes from wild-type (red) or *CCAR1* CRISPRi knockdown K562 cells (blue) (more examples are shown in **Figure S6**). On the right, each noted exon inclusion event was validated using specific primers (see **Table 6**) for the corresponding sites in *CCAR1* wild-type and knockdown samples. In all cases, the lower band represents the wild-type splicing pattern, while the upper band represents the exon-included version. **D)** Illustration of CCAR-dependent alternative splicing events in mouse cells. Sashimi plots display CCAR-dependent exon usage in the noted genes from wild-type (red) or *CCAR1^-/-^* 3T3 cells (blue) (more examples are shown in **Figure S6**).

Using RT-PCR, we validated several of the RNA-seq identified CCAR1-dependent exons (**Figure 4C, S8**). In all cases, the relevant cassette exons were included or excluded from the mRNA upon *CCAR1* knockout. This includes a known and conserved PE in *IVNS1ABP*, which we now identify as CCAR1-dependent (Manet, 2021). Other interesting CCAR1-dependent splice events include a novel transcription start site for *ITPK1*; novel internal exons for *Heatr5a, ITPK1, and WSB1*; and alternative splicing in multiple disease-related genes such as *CD34, Senp6* and *Max* (**Figure 4D**). The functional consequences of these alternative splicing events remain to be determined, but many of them alter coding sequence.

*FANCA* was not among the differentially spliced mouse transcripts, but we noted that the *FANCA* PE is a relatively recent evolutionary development that is only found in primates (**Figure S7B**). On a global level, only some of the CCAR1-dependent splicing changes are concordant between human and mouse (**Figure S7C)**. However, CCAR1’s role in regulating splicing is evolutionarily conserved, and it plays a key role in regulating many splicing events.

## Discussion

Utilizing a comprehensive genome wide CRISPRi screening strategy, we searched for novel regulators of CRISPR-Cas mediated HDR. We surprisingly found that *CCAR1* plays a critical role in supporting HDR and interstrand crosslink repair by ensuring normal splicing of *FANCA*. The cellular function of *CCAR1* is poorly understood, but it is mostly discussed as a transcriptional regulator ^14,15^. We found that CCAR1 unexpectedly controls multiple splicing events in human and mouse cells, mostly regulating the incorporation of cassette exons. This is consistent with isolated reports that *CCAR1* regulates alternative splicing of certain neuronal genes in mouse ^32^ and unc-52/perlecan in *C.elegans* epidermis ^33^. Our findings reveal a widespread and conserved role of CCAR1 in shaping alternative splicing across species.

CCAR1’s role in splicing could be direct or indirect. Publicly available proteomic databases indicate that CCAR1 has unappreciated interactions with multiple components of the spliceosome, including U2AF2 (**Figure 5A**)^34^. CCAR1 has also been detected in unbiased yeast-two-hybrid experiments as potentially part of the spliceosome A complex, which is recruited to the exon-intron junctions at the early splicing cycle ^34,35^, ENCODE eCLIP datasets in wild type cells indicate that the U2AF1 and U2AF2 core splicing factors bind to many of the CCAR1-regulated exons we identified in our unbiased RNA-seq datasets, even though these exons are not normally spliced in a wild type background (**Table 5, Figure S9**). This implies that CCAR1 may be directly repressing exon inclusion, despite spliceosome engagement. Of note, U2AF1 and U2AF2 are the two top splicing factors whose perturbation affected *FANCA* PE inclusion in ENCODE data (**Figure S4**). On the other hand, knockdown of *CCAR1* in both human and mouse cells also consistently transcriptionally downregulated RNA binding proteins and splicing factors (**Figure 5B**). Thus, we propose that CCAR1’s widespread role in regulating splicing is due to a direct interaction with and modulation of spliceosome activity, and/or indirect regulation by altering the abundance of splicing factors (**Figure 5C**).

**Figure 5.**
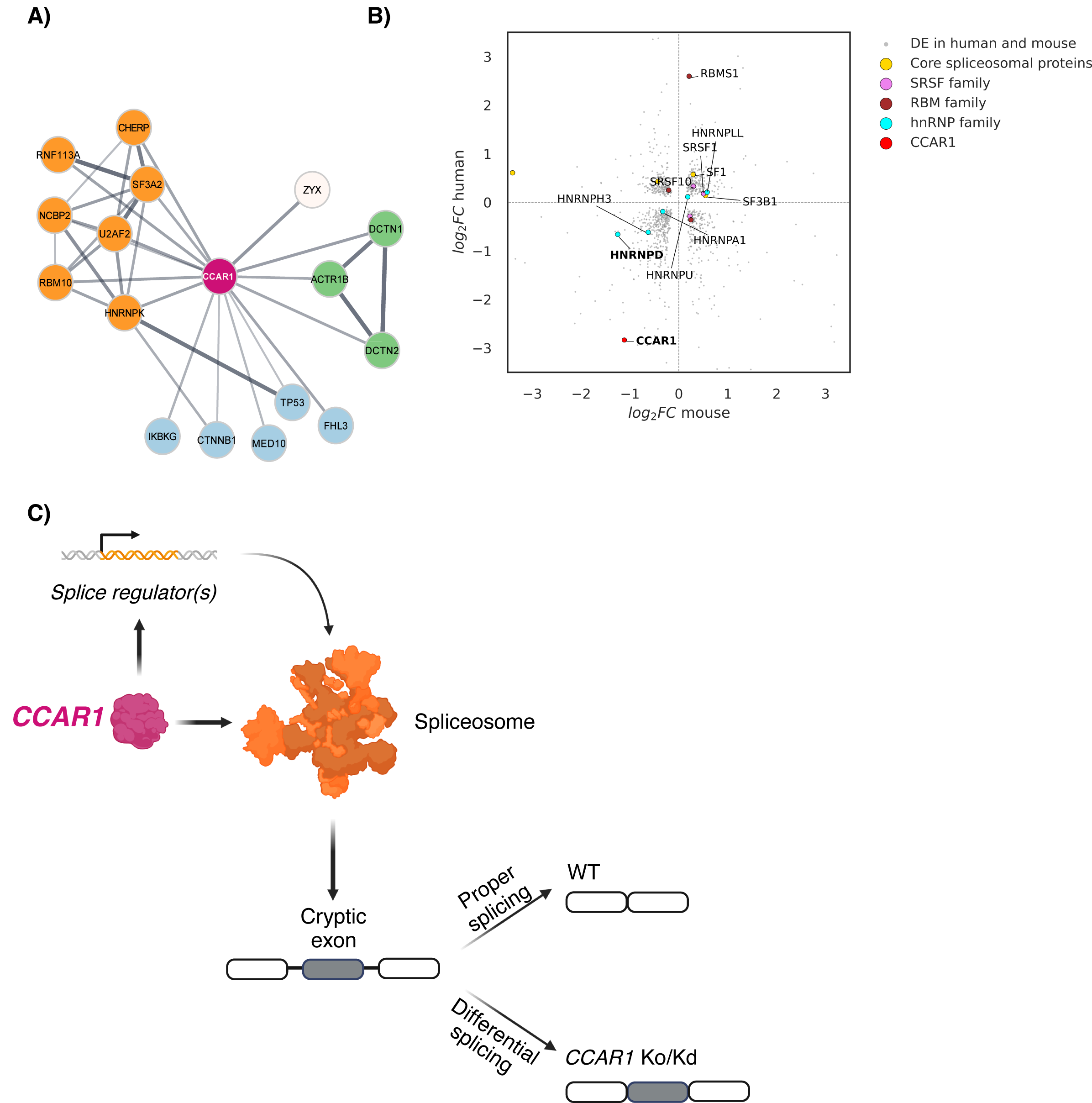
CCAR1 may act as a fidelity factor for spliceosome. **A)** CCAR1 interacts with transcription machinery (blue) and multiple spliceosome components (orange) in the STRING database. **B)** Knockdown of CCAR1 leads to consistent transcriptional dysregulation of select HNRNP splicing factors in both of our mouse and human RNA-seq datasets. **C)** Model of CCAR1-mediated regulation of cryptic exon inclusion. *CCAR1* directly interacts with the spliceosome to prevent poison exon inclusion and/or regulate the abundance of RNA binding proteins or splicing factors to maintain splicing fidelity. Illustrations were created with BioRender.

To the best of our knowledge, *the FANCA* PE is the first poison exon identified in a DNA repair gene. Given the potent loss of FANCA protein expression when CCAR1 is absent, one might hypothesize that CCAR1 could represent a missing FA complementation group. That is, human patients with loss of CCAR1 activity might display FA-like phenotypes. However, no CCAR1 mutations have been reported in the FA mutation database (https://www2.rockefeller.edu/fanconi/), and we are unaware of any reports of FA patients carrying CCAR1 mutations. Instead, CCAR1 loss of function variants are almost completely missing from the human population (gnOMAD pLOEUF = 0.21) ^36^. Given CCAR1’s widespread roles in regulating transcription and splicing, we propose that non-functional CCAR1 mutations are inimical to development.

Poison exons are generally highly conserved ^37^ and functional PEs are primarily found within genes encoding RBPs, forming dense post-transcriptional expression regulatory networks ^38^. Moreover, deregulation of the control of PEs is associated to neurological diseases such as Alzheimer’s, Parkinson and dementia ^38–40^. Although the most affected exon in *CCAR1* knockdown cells was the FANCA PE, this exon seems to have emerged relatively recently during primate evolution. It is currently unclear how CCAR1 developed the ability to regulate this evolutionarily recent PE in human cells, as well as whether CCAR1 uses the same or distinct molecular partners to impact exon inclusion in human and mouse cells. We found no obvious conserved sequence motifs upstream or downstream of CCAR1-responsive exons, but splice regulatory sequences are typically short and difficult to detect from primary sequence alone (**Figure 4C, D**). The exact context that differentiates CCAR1-responsive from CCAR1-insensitive exons remains to be determined.

## Supporting information

table1_drugZ_analysis

table1_mageck_analysis

table2_de_human_list

table3_human_diff_spliced_junctions

table4_human_diff_spliced_junctions

table5_ENCODE_eclip_analysis

table6_primer_list

## Data Availability

Sequencing files for the pooled screen and RNA-seq datasets will be uploaded to GEO/SRA.

## Ethics declarations

### Code Availability

The code used to analyze CRISPR screen and RNA-seq would be available upon request.

### Author Contributions

M.E.K and J.E.C conceived the study. M.E.K designed and performed wet-lab experiments. Analysis of RNA-seq for deregulated genes and alternative splicing is done by B.J and A.M. A.M has produced the figure5 B and data for figure 2 C. B.J generated the graphs and pie charts for Figure4 A and B. M.E.K, A.G and J.E.C wrote the first draft of the paper with input from other authors.

## Acknowledgements

We thank the Functional Genomics Center Zurich (FGCZ) and especially Dr. Susanne Kreutzer and Dr. Zacharias Kontarakis for their help with CRISPR screen NGS sequencing and RNA-seq experiments. We thank Dr. Chaolin Zhang for kindly providing access to the serves of his lab and help to run the alternative splicing analysis. We also thank the members of the Corn Lab for helpful discussions and help with the manuscript.

JEC is supported by the NOMIS Foundation and the Lotte and Adolf Hotz-Sprenger Stiftung. MEK is supported by the Fanconi Anemia Research Foundation. This project has received funding from the European Research Council (ERC) under the European Union’s Horizon 2020 research and innovation programme (grant agreement No 855741, DDREAMM). This work was supported as a part of NCCR RNA & Disease, a National Centre of Competence (or Excellence) in Research, funded by the Swiss National Science Foundation (grant number 205601).

## METHODS

### Cell lines

K562 cells were cultured in RPMI 1640 GlutaMAX™ medium supplemented with 10% FBS and 1% P/S solution. RPE1 cells were cultured in DMEM, high glucose, GlutaMAX™ pyruvate medium (ThermoFisher Scientific, 10569010) supplemented with 10% FBS and 1% P/S. All cells were maintained at 37°C with 5% CO2 in a humidified incubator. Regular mycoplasma testing was performed using the MycoAlert Mycoplasma Detection Kit (Lonza, LT07-318). To generate RPE1 CRISPRi cell lines, the CRISPR inhibition plasmid (pHR-EF1a-dCas9-HA-mCherry-KRAB-NLS) construct was incorporated into lentiviruses in HEK293T cells. The lentiviral supernatant was filtered and then used to transduce RPE1 cells. After transduction, mCherry-positive cells were isolated by sorting using the SH800 Cell sorter (Sony). K562 CRISPRi (K1e) cells were previously generated in a prior study ^11^.

### CCAR1 knock out K562 human and 3T3 mouse cell line generation

The gRNAs were synthesized in vitro as per the described procedure ^45^. In brief, overlapping oligomers containing a T7 promoter, protospacer, and gRNA scaffold, as listed in Table 6, were subjected to amplification using Q5 High-Fidelity polymerase (New England Biolabs, M0491L) for 15 cycles. A mixture of 1 µM T7FwdLong and T7RevLong served as the template and was amplified by T7FwdAmp and T7RevAmp in a 50 µl reaction volume. Subsequently, 8 µl of the PCR-amplified product was employed for in vitro transcription using the NEB HiScribe T7 High Yield RNA Synthesis kit (New England Biolabs, E2040S), with incubation at 37 °C for 18 hours in a thermocycler. The reaction was then supplemented with DNase I (Qiagen, 79256) for 30 minutes at 37 °C, followed by treatment with Quick CIP (New England Biolabs, M0525S) for 1 hour at 37 °C. The resulting gRNAs were purified using the miRNeasy kit (Qiagen, 217604), their concentration was quantified using the Qubit™ RNA Broad Range (BR) assay (ThermoFisher Scientific, Q10211), and they were stored at −80 °C. To target the *CCAR1* locus, a mixture containing 100 pmol SpCas9-NLS and 120 pmol gRNA in Cas9 buffer was prepared and incubated for 20-30 minutes at either room temperature or 37°C. A cell count ranging from 1 × 10^5 to 2 × 10^5 cells was collected and suspended in 15 µL of nucleofection buffer (Lonza). Electroporations were conducted in the strip format, utilizing a total volume of 20 µL that included both cells and the RNP mix. The choice of kit and program was determined based on the specific cell type: K-562 cells were electroporated using the SF kit and the FF-120 program, while 3T3 cells were subjected to electroporation using the SG kit and the EN-158 program. After electroporation, 80 µL of prewarmed DMEM or RPMI medium was promptly added to the strips. The cells were allowed to incubate in a biosafety hood for 10 minutes before being transferred to culture plates and returned to a 37°C incubator. Following a recovery period of 2-3 days, the cells were seeded in 96-well plates as single cells, and colonies were subsequently analyzed either by Next-Generation Sequencing (NGS) or Sanger Sequencing to assess *CCAR1* knockout alleles.

### Sanger and next generation (NGS) sequencing

Genomic DNA extraction was performed using QuickExtract™ DNA Extraction Solution (Lucigen), and the genomic locus of interest was amplified using AmpliTaq Gold® 360 Master Mix (ThermoFisher Scientific). For NGS library preparation, two rounds of PCR were conducted. In the first round (PCR 1), the PCR primers contained sequences corresponding to the genomic locus, along with the appropriate forward and reverse Illumina adapters. PCR 2 was carried out using unique Illumina barcoding primer combinations, with 1.5 µl of amplified product from PCR1.

The PCR2 products were purified using SPRIselect beads (Beckman), with a beads-to-PCR product volume ratio of 0.9×. Prior to multiplexing, the resulting amplicon size and concentration were verified on the 4200 TapeStation System (Agilent). For Sanger sequencing (and PCR1 for NGS), the PCR products were purified using MinElute columns (Qiagen) and eluted in 30 µl elution buffer (EB). Approximately 120 ng of purified PCR product was then submitted for Sanger economy sequencing (Microsynth). Detailed primer information can be found in Table 6.

### Plasmid generation for CRISPRi guides

The sgRNAs were designed to include a variable 20-nucleotide sequence that corresponds to the target gene. Oligos for both sgRNAs and nicking guides were ordered from IDT and subsequently cloned into the pLG1-puro-BFP vector, which contains dimBFP, after digestion with BstXI and BlpI enzymes.

### Total RNA extraction and cDNA generation

RNA extraction was conducted using the RNeasy Mini Kit from Qiagen, following the manufacturer’s instructions. For each sample, 1 μg of total RNA was utilized for reverse transcription, employing the iScript™ Reverse Transcription Supermix to generate complementary DNAs (cDNAs).

### Plasmid generation for CCAR1 fragments and FANCA

CCAR1 open reading frames (ORFs) were generated through PCR amplifications using K562 cDNA as the source. These ORFs were subsequently cloned into pLenti-X1 destination vectors, which carried either neomycin or puromycin resistance markers. The pLenti-X1 vector was initially digested with the restriction enzymes BamHI and XbaI. Subsequently, Gibson assembly was employed to insert the gene of interest along with small protein epitopes into the pLenti-X1 vector ^46^. BioAD tag ^47^ construct was kindly gifted by Dr. Stefanie Jonas, ETH Zurich.

For FANCA cDNA, the FANCA ORF was transferred from a vector (phEF1a-FANCA-IRES-Thy1.1) ^11^ into pLenti-X1 vectors containing neomycin resistance. In brief, the FANCA ORF was amplified from the phEF1 vector, and Gibson assembly was utilized to insert it into the pLenti-X1 vector.

### Plasmid generation for Mini-gene reporters

The specified FANCA genomic region was amplified from K562 genomic DNA. The pcDNA3 vector was linearized through the BamHI and EcoRI restriction sites. Subsequently, Gibson assembly was employed to insert the amplified FANCA regions into the pcDNA3 vector.

For scrambling mutagenesis, random sequence scrambling was performed, and the fragments with the mutated sequences were amplified from the poison exon-positive (PE+) mini gene construct. These fragments were later inserted into the pcDNA3 vector using Gibson assembly.

### Western blotting and MMC treatment

For MMC treatment, 10^6^ K562 cells were incubated with 1 µM MMC for 24 h. For protein extracts, 1 × 10^6^ LCLs were pelleted and washed in PBS. and washed with PBS. The cells were then lysed in ice-cold RIPA buffer (Millipore) supplemented with Halt protease inhibitor cocktail (ThermoFisher Scientific) and Benzonase (200 Units), unless otherwise specified. After centrifugation at 21,000 × g for 30 minutes at 4 °C, the supernatant was transferred to another microcentrifuge tube. Protein concentration was determined using the Bradford Assay (VWR). Protein lysates were diluted in RIPA buffer supplemented with 1× LDS and 1× DTT and heated at 95 °C for 5 minutes. Approximately 15-30 total µg of protein was loaded onto 4–12% polyacrylamide gels (NuPAGE) with 1× MOPS SDS running buffer (NuPAGE).

Proteins were transferred using the Criterion Trans-Blot® Cell (BioRad) with Tris-Glycine transfer buffer (25 mM Tris base, 192 mM glycine, 20% methanol (v/v); pH = 8.3). The membrane was briefly stained with Ponceau staining to confirm the transfer and then blocked with 5% (w/v) non-fat dry milk in TBS-T (0.1% Tween-20) for 1 hour at room temperature. Primary antibodies, including anti-rabbit FANCA antibody (A301-980A-T, Thermo Scientific and ab5036, Abcam), anti-rabbit FANCD2 antibody (ab221932, Abcam), anti-goat HSP60 antibody (sc-1052, Santa Cruz Biotechnology), anti-rabbit GAPDH (2118S, Cell Signaling), or anti-rabbit FLAG (14793S, Cell Signaling), were diluted 1:1000 in 5% BSA-TBS-T. HSP60 or GAPDH antibodies were used as loading controls. The membrane was incubated with primary antibodies overnight at 4 °C and then washed three times with TBS-T. Subsequently, the membrane was incubated with anti-rabbit secondary antibody (IRDye 800CW (926-32213) or anti-goat IRDye 800CW (926-32214)) at a 1:5000 dilution in 10% milk TBS-T for 45 minutes.

Finally, the membrane was washed twice with TBS-T and once with PBS before being imaged using Li-Cor’s Near-InfraRed fluorescence Odyssey CLx Imaging System.

### MMC sensitivity assay

The MMC sensitivity assay was conducted in two different manners. In Figure 1C, K562 cells were plated in 96-well plates at a density of 1000 cells per well. They were then exposed to increasing concentrations of MMC (0, 3, 10, 33, 100, 333, 1000 nM). Cell survival was assessed using the CellTiter Glo reagent.

In Figure S2, cells transfected with the specified guide RNAs were mixed with wild-type cells that did not express any fluorophore. This mixed cell population was then split into two separate wells. One well served as a control, while the other well was incubated with a lower dose of MMC (50 nM) for a duration of 5 days. After this incubation period, the cell mixtures were analyzed using flow cytometry, and downstream analysis was conducted using FlowJo Software v10.7.1 (FlowJo, LLC) to measure the BFP (blue fluorescent protein) ratio within the population.

### CRISRi Screen in K562 cells

#### Genome-wide library construction and lentivirus production

As previously mentioned in ^13^, the genome-wide CRISPRi-v2 library ^48^ was transferred into a lentiviral vector designed for the expression of guide RNAs. This lentiviral vector also contains a blue fluorescent protein (BFP) marker. The lentiviruses carrying this library were produced using HEK293T cells as packaging cells.

#### CRISPR screen

The K562 CRISPRi cells were cultured and grown until they reached a population size of approximately 150 million cells. Subsequently, they were transduced with the genome-wide CRISPRi library at a multiplicity of infection (MOI) of 0.3. The transduction efficiency was monitored by identifying BFP-positive cells, and the coverage reached approximately 500-fold for each individual guide RNA (sgRNA).

Two days after transduction, the cells containing the guide RNAs were subjected to puromycin selection at a concentration of 2 µg/ml. This selection process continued for 96 hours to ensure the survival of cells expressing the guide RNAs. After successful puromycin selection, the cell population was divided into two replicates, and was maintained a coverage of 500-fold for each sgRNA throughout the screening process.

A week after the transduction, 20 × 10^6 was subjected to electroporation. This involved the introduction of 400 pmol of SpCas9-NLS, 480 pmol of L2 guide RNA (gRNA) targeting BFP, and 500 pmol of a single-stranded oligodeoxynucleotide (ssODN) template containing the sequence for converting BFP to GFP. Electroporation was carried out using FF120 and SF solution in a 4D electroporator. This process was performed twice for each replicate, totaling six electroporations.

Following electroporation, the cells were expanded in culture before undergoing sorting. Prior to sorting, a background population of cells was collected for subsequent next-generation sequencing (NGS) analysis. The sorting step was aimed at enriching populations of cells that were either GFP-positive or GFP-negative. After sorting, the cells were pelleted and stored at −80°C until genomic DNA extraction for further analysis.

#### NGS sample preparation and screen analysis

Genomic DNA was extracted using Gentra PureGene Cell Kit (Qiagen, 158912) gDNA extraction protocol.

Briefly, cell pellets were first resuspended in 3 ml of cell lysis solution and then mixed with 15 µl of RNase A solution. This mixture was incubated at 37°C for 20 minutes, followed by a cooling step on ice for 10 minutes. Subsequently, 1 ml of protein precipitation buffer was added to the mix, which was vortexed thoroughly. Afterward, the mixture was centrifuged at 2000 × g for 10 minutes to pellet cellular debris. The supernatant, which contained the genomic DNA, was then mixed with 3 ml of 100% isopropanol by gently inverting the 15 ml tube approximately 50 times. Genomic DNA was pelleted by centrifugation at 2000 × g for 5 minutes and then washed with 70% ethanol (EtOH). Following the ethanol wash, any remaining 70% EtOH was removed, and the genomic DNA was resuspended in 200 µl of hybridization buffer. The resuspended DNA was incubated at 50°C for 1 hour, and the DNA concentration was subsequently measured using a Nanodrop instrument.

Purified genomic DNA were used for the further PCR amplification of sgRNA cassettes following the protocol (<u>weissman.wi.mit.edu/resources/IlluminaSequencingSamplePrep.pdf</u>). For the background samples, 5 µg of genomic DNA was used per reaction. To prepare the sort background samples, 30 PCR reactions were initially performed and then combined. In the case of GFP-positive cells, 10 PCR reactions were conducted and later pooled, while for GFP-negative cells, 15 PCR reactions were performed and subsequently combined. The PCR products underwent purification through two rounds of Sera-Mag magnetic beads (Cytiva, catalog number 29343957). The concentrations of the samples were determined using the Qubit™ 1× dsDNA high-sensitivity assay (ThermoFisher Scientific, catalog number Q33232), and the samples were grouped together based on their expected read counts. Subsequently, the samples were subjected to sequencing on the NextSeq 2000 platform.

Guide RNA abundances within each population were assessed through targeted next-generation sequencing. Subsequently, MAGeCK ^49^ and DrugZ ^50^ analysis tools were employed to translate guide RNA abundances into gene-level phenotypes and determine statistical significance, as depicted in **Figure 1A** and summarized in **Table 1**.

#### RNP electroporation for BFP to GFP reporter assay

In summary, 36 pmol of sgRNA and 30 pmol of SpCas9-NLS were combined in Cas9 buffer (containing 20 mM HEPES at pH 7.5, 150 mM KCl, 1 mM MgCl2, 10% glycerol, and 1 mM tris(2-carboxyethyl)phosphine (TCEP) reducing agent). This mixture was allowed to incubate at room temperature for 20 minutes. Simultaneously, 1 × 10^5 to 2 × 10^5 cells were harvested and centrifuged at 300 × g for 5 minutes. The resulting cell pellets were resuspended in 15 µL of nucleofection buffer (Lonza). Subsequently, 5 µl of the RNP mixture was added to the cell suspension, along with 0.3 µl of a 100 µM (30 pmol) ssODN (BFP to GFP template). For RPE1 cells, phosphorothioate bond-containing ssODN templates were used, as indicated in ^13^. Five days after electroporation, the cells were collected and subjected to flow cytometry analysis using an Attune Flow Cytometer (ThermoFisher Scientific). The downstream analysis was carried out using FlowJo Software v10.8.2 (FlowJo, LLC).

Electroporations were performed in the strip format, with 20 µl volume of cells and RNP mix. The following kit and program for each cell type was selected: K-562 (SF kit/FF-120), RPE1 (P3 kit/ EA-104). After electroporation, prewarmed 80 µl of DMEM or RPMI medium was added into strips. Cells were incubated in the hood for 10 mins and then transferred to the plates and returned to 37 °C.

### Quantitative real-time PCR (qRT-PCR)

RNA extraction was carried out utilizing the RNeasy Mini Kit (Qiagen), following the manufacturer’s prescribed protocol. For each sample, 1 μg of RNA was utilized for reverse transcription, which was conducted using the iScript™ Reverse Transcription Supermix for RT-qPCR (Biorad, 1708841), following the manufacturer’s instructions.

The qRT-PCR reactions were prepared using SsoAdvanced Universal SYBR Green Supermix (Biorad, 1725271) and were performed in triplicate runs. The QuantStudio 6 system (ThermoFisher Scientific) was employed for running the qRT-PCR experiments. A comprehensive list of primers employed in the RT-qPCR can be found in Table 6.

#### Mini-gene reporter assays and endogenous FANCA PE inclusion RT

pcDNA vectors containing mini-gene reporters, approximately 1 μg in quantity, were introduced into both wild-type and *CCAR1*-suppressed K562 cells via electroporation. Following a 2-day incubation period, K562 cells were harvested, and total mRNA was extracted. Subsequently, cDNA was synthesized as previously described. The primers used in this process targeted the constant regions of the vectors to amplify the spliced mRNA products (see table 6).

To identify the presence of endogenous FANCA poison exon (PE), cDNAs from the specified samples were employed along with primers that bound to exon 14 and exon 15, respectively. A brief PCR was conducted to amplify the region of interest, and 2.5% agarose gels were prepared to facilitate the separation of the PE-included and wild-type PCR-amplified bands

#### RNA-seq library prep

K562 and 3T3 cells were cultured, harvested, and subjected to RNA extraction using the RNeasy Mini kit (product number 74104) along with the RNase-Free DNaseI Kit (product number 79254). The concentrations of RNA were quantified using the Qubit RNA BR Assay from ThermoFisher, and 1 µg of total RNA was utilized for library preparation. For RNA-seq library construction, the Illumina TruSeq RNA Library Prep Kit v2 was employed, following the manufacturer’s instructions. In the case of K562 cells, three replicates were prepared for *CCAR1* knockdown samples, as well as for wild-type K562 cells. These libraries were subsequently sequenced on a NextSeq 2000 instrument, generating 150 paired-end reads.

For 3T3 cells, three replicates were prepared for *CCAR1* knockout samples, and two replicates for wild-type 3T3 cells. These libraries were sequenced on a NextSeq 2000 instrument, producing 50 paired-end reads as the sequencing output.

#### RNA-seq analysis

Raw sequencing data was mapped to the Human reference genome sequence (GRCh38.p13) or mouse reference genome sequence (GRCm38.p6) using default settings on HISAT2 ^51^. Gene counts were obtained by assigning and counting reads to the human and mouse gene annotations (GENCODE Release 36 – GRCh38.p13, GENCODE Release M2, GRCm38.p6) using Rsubread ^52^. Differential gene expression analysis was performed using the R package DESeq2 ^53^ and applying a strict adjusted *p*-value cutoff of 0.05.

#### Splicing analyses

Splice junctions were mapped using the MAJIQ algorithm (2.0) under default conditions ^31^. Splice graphs and known/novel local splice variants (LSVs) were defined through use of the MAJIQ builder provided with comprehensive human and mouse gene annotations (GENCODE Release 36 – GRCh38.p13, GENCODE Release M2, GRCm38.p6) and uniquely mapped BAM files as input. The MAJIQ Quantifier was used to calculate relative abundances (Percent selected index – PSI) for all defined junctions from the perspective of sources and targets ^31^. Following this, the MAJIQ dPSI function was used to compare different conditions and the Voila “modulizer” method (--decomplexify-reads-threshold 10 --changing-between-group-dpsi 0.1) was used to detect specific classes of binary splicing events, and output those that underwent significant changes between specified conditions into tabular formats.

Custom R scripts were written to process Voila outputs to obtain significantly changing splice junctions. Briefly, we used probability_changing > 0.8 (confidence level) and mean_dpsi_per_lsv_junction > 0.1. Together, these criteria provided junctions with strong statistical likelihood of at least 10% change between conditions. Furthermore, after manually vetting a substantial number of events, we adopted the classifications attributed by MAJIQ to each splice junction (such as cassette exon, alternative 3’SS *et cetera*). MAJIQ also defined *de novo* junctions based on the provided GENCODE annotations. We adopted a more stringent definition in which *de novo* junctions were also required to have a PSI < 0.05 in control conditions. This allowed us to distinguish *de novo* splicing that was due to poor annotation of human or mouse genes, versus those that are likely to be new splice junctions. Sashimi plots were generated using the Integrative Genomics Viewer (IGV) ^54^.

#### FANCA poison exon inclusion analysis

The database of rMATS- and Deseq2-processed RNA-binding protein (RBP) perturbations available at the ENCODE portal ^43^ and SRA archive, was constructed before ^55^. The database was queried for the rMATS output for poison exon 15 within the FANCA gene. The log2FC values of the targeted proteins were obtained from the same database by querying for the Deseq2 output.

#### RNA-seq data to get differentially expressed genes

Paired-end reads from the raw fastq files of human and mouse samples were mapped to human genome version hg38 and mouse genome version mm39, respectively, using STAR v. 2.7.10b ^56^. The following parameters were used: --outFilterType BySJout --outSJfilterOverhangMin 10 10 10 10 --alignSJDBoverhangMin 10 --alignSJoverhangMin 10 --outFilterMismatchNmax 999 --outFilterMismatchNoverReadLmax 0.05 --outFilterScoreMinOverLread 0.4 --outFilterMatchNminOverLread 0.4 --twopassMode Basic --outFilterMultimapNmax 500000000. GENCODE ^57^ comprehensive annotations v42 and vM32 were used for human and mouse samples, respectively. RNA-SeQC ^58^ was utilized to obtain raw gene counts from uniquely mapped exonic reads. Differential expression (DE) analysis was performed using the Deseq2 ^53^ R package version 1.40.2 with default parameters.

---

**Figure S1.**
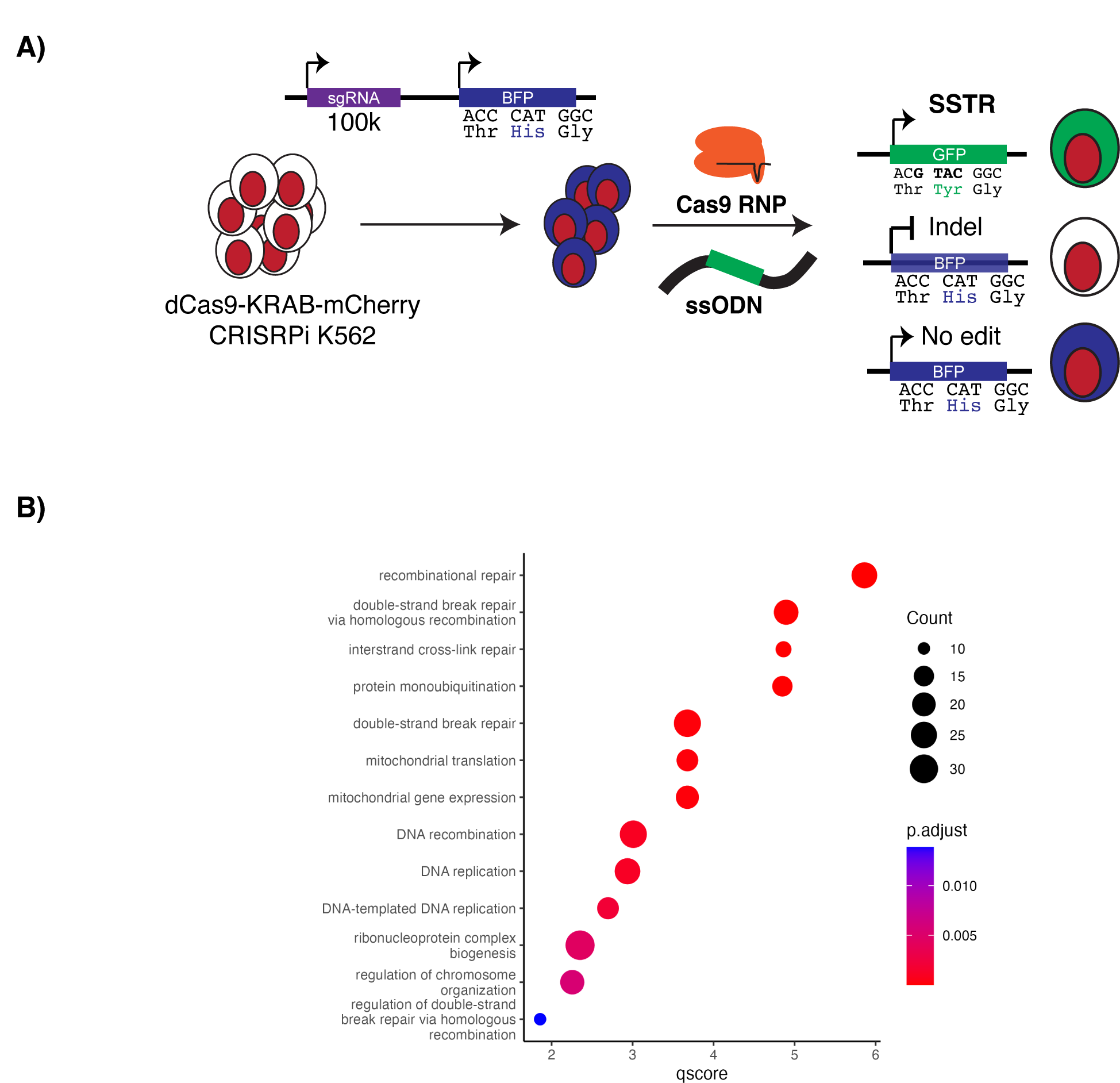
Genome-wide CRISPRi screen interrogating novel HDR players. **A)** The diagram illustrates the genome-wide screening strategy, explained in detail in the Methods section. **B)** Unbiased Gene Ontology (GO) ontology analysis of significant hits revealed that genes depleted from the GFP-positive population (MAGeCK score <= 0.01 and DRUGZ normZ <= −0.5) are significantly enriched in the double-strand break repair via homologous recombination and the Fanconi Anemia repair pathway. GO-term analysis was performed using GOSemSim ^41^ and clusterProfiler ^42^.

**Figure S2.**
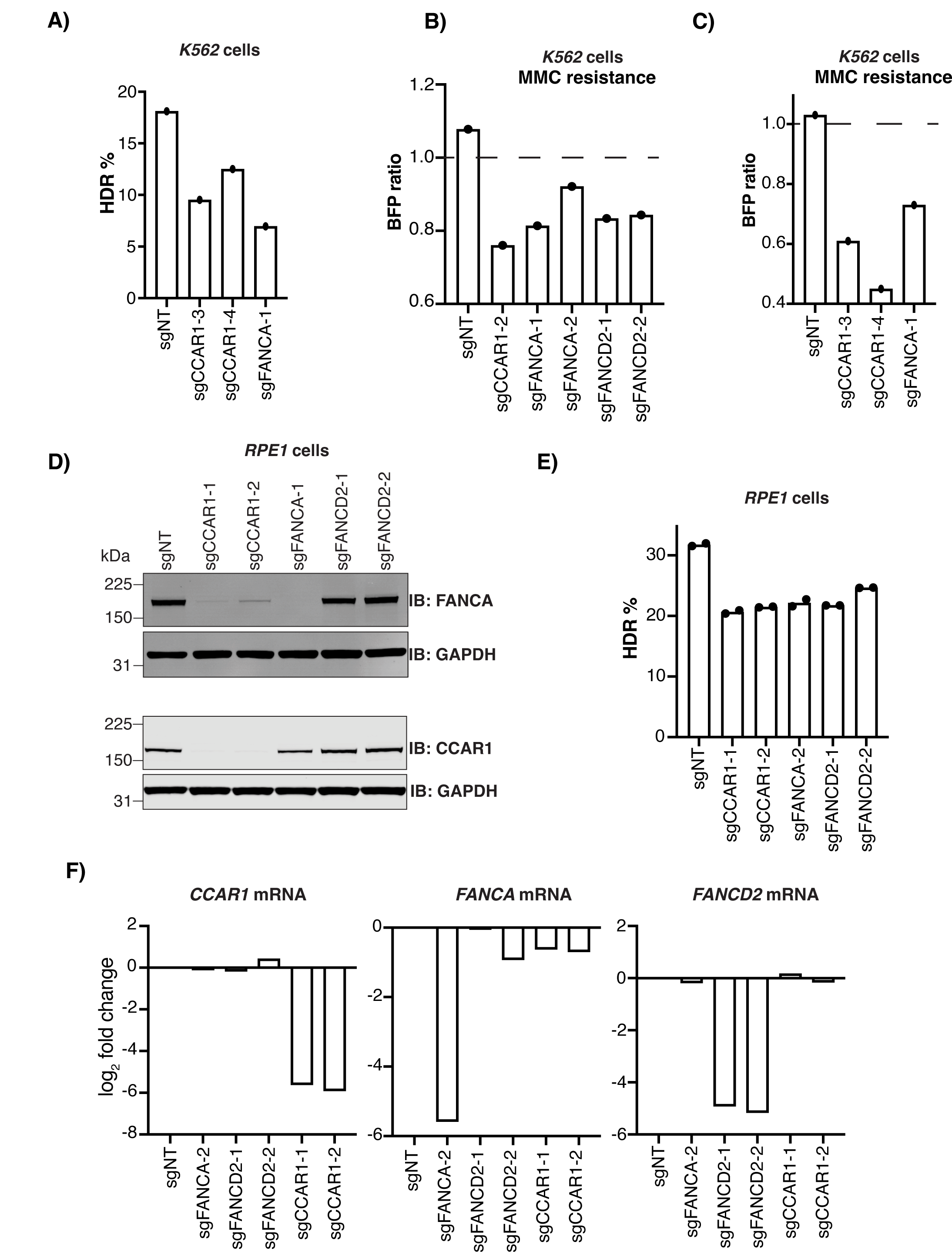
Validation of FA deficiency phenotype in *CCAR1* depleted cells. **A)** Two additional sgRNAs targeting *CCAR1* were used in the BFP-GFP HDR assay, as described in Figure 1C. Each dot represents individual biological replicate, and the bars represent the mean. **B, C)** MMC sensitivity was assessed in a competition assay in K562 cells. Stably expressing CRISPRi clones (BFP positive) were mixed with wild-type (no fluorophore) cells and incubated with or without 50 nM MMC for 5-days. Cells were then analyzed by flow cytometry and the ratio of BFP-positive vs. BFP-negative cells was measured. **D)** FANCA protein is abolished in the absence of *CCAR1* in RPE1 stable CRISPRi cells. Protein extracts were analyzed by Western blot using the indicated antibodies (anti-FANCA, anti-CCAR1, anti-GAPDH). FANCD2 was detected as a doublet, and the slowly migrating FANCD2 band represents FANCD2-Ub. The lower panel displays the Western blots from *CCAR1* knockdown RPE and wild-type cells probed with anti-FANCA and anti-CCAR1 antibodies. **E) Stable** CRISPRi knockdown of *CCAR1* reduces CRISPR-Cas9 mediated HDR in RPE1 cells. RPE1 CRISPRi cell lines were stably transduced with individual sgRNAs targeting *CCAR1*, *FANCA*, and *FANCD2*. The BFP-to-GFP assay was used to measure HDR efficiency. Each dot represents individual biological replicate, and the bars represent the mean. **F)** FANCA mRNA levels show minimal changes in RPE1 cells. RT-qPCR analysis of RNAs extracted from the indicated cell lines. The plotted values represent the log_2_ fold difference normalized to the healthy donor sample. Bars indicate means for each cell line.

**Figure S3.**
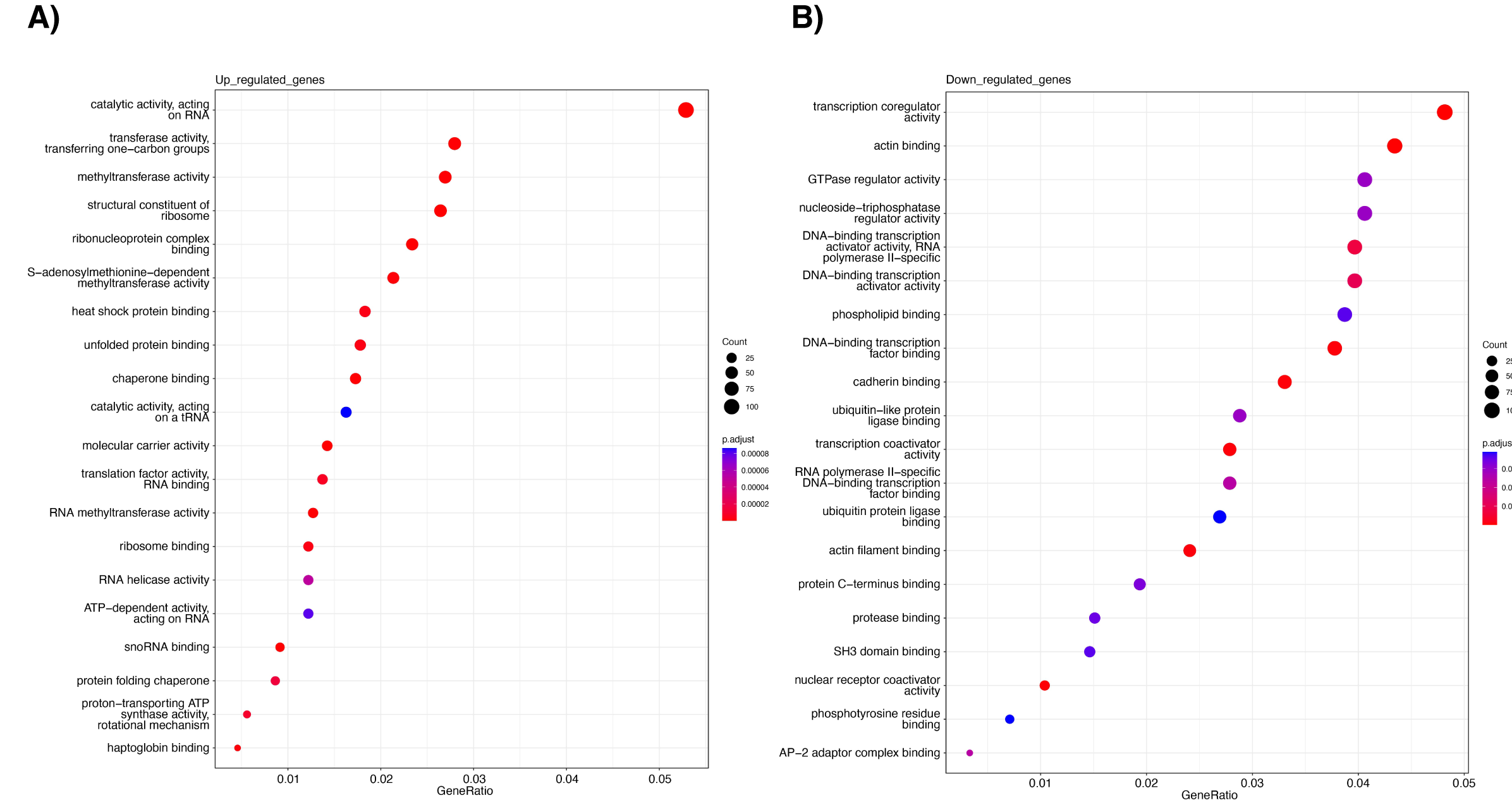
Unbiased gene ontology analysis in the K562 RNA-seq cell lines. Significantly up regulated (**panel A**) and down regulated (**panel B**) genes observed in CCAR1-knockdown cells were summarized with GO-term analysis as explained in the Methods section.

**Figure S4.**
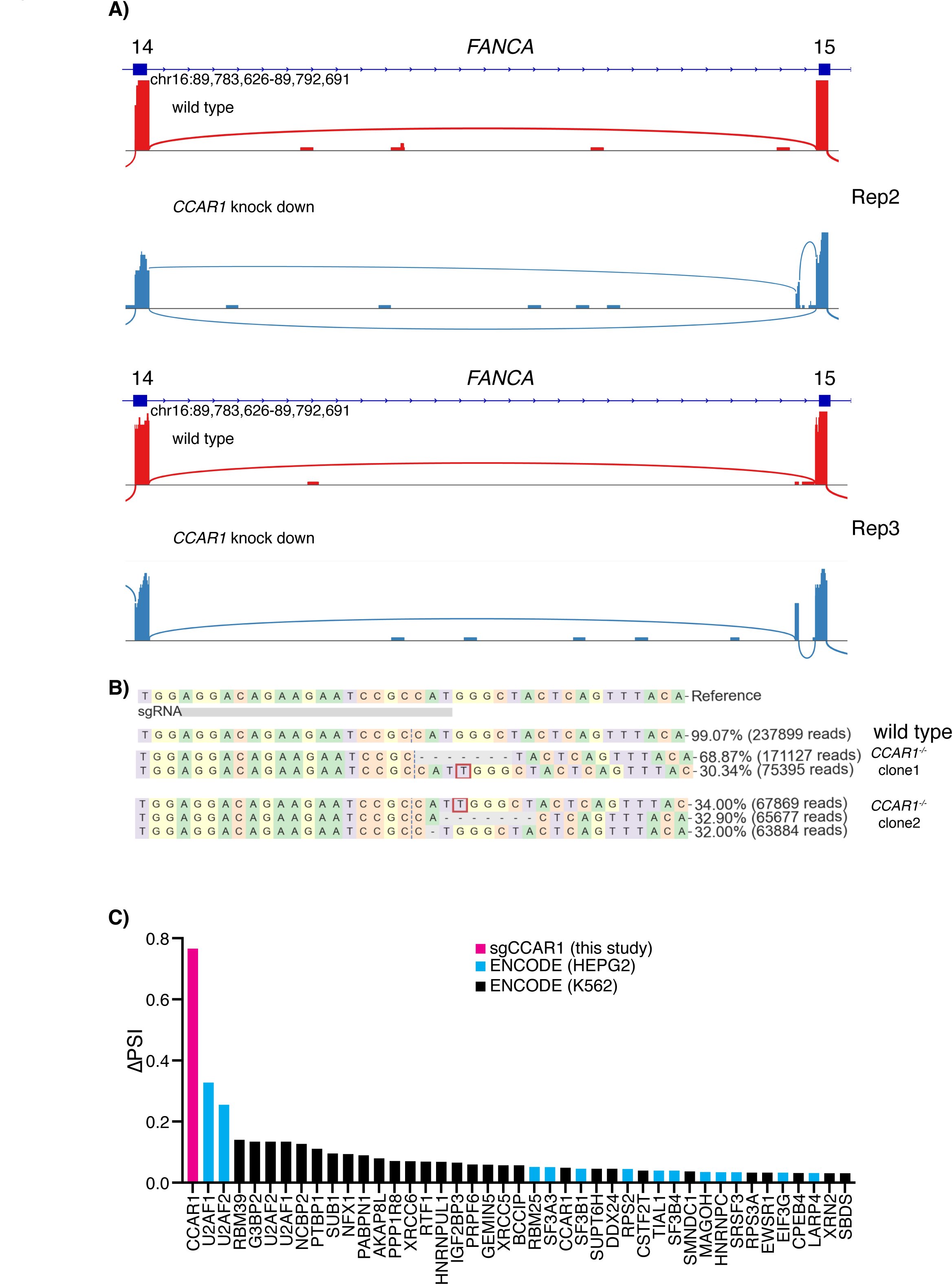
FANCA PE is present in the ENCODE datasets, albeit at lower levels. **A)** Sashimi plots of FANCA PE exon inclusion from replicates 2 and 3 of the human RNA-seq dataset. **B)** Next-generation sequencing confirms the human *CCAR1^-/-^* K562 knock out clones. CRISPResso2 output shows the compound heterozygous deletions in the *CCAR1* locus. **C)** Re-analysis of differential exon inclusion in splice factor perturbation datasets from ENCODE project ^43^. Several RNA binding proteins were suppressed in K562 or in HEPG2. As a comparison, CCAR1 depletion from this study was added in the graphs (magenta bar). ΔPSI values calculated from different datasets for the FANCA PE event are visualized in the graphs. See Methods.

**Figure S5.**
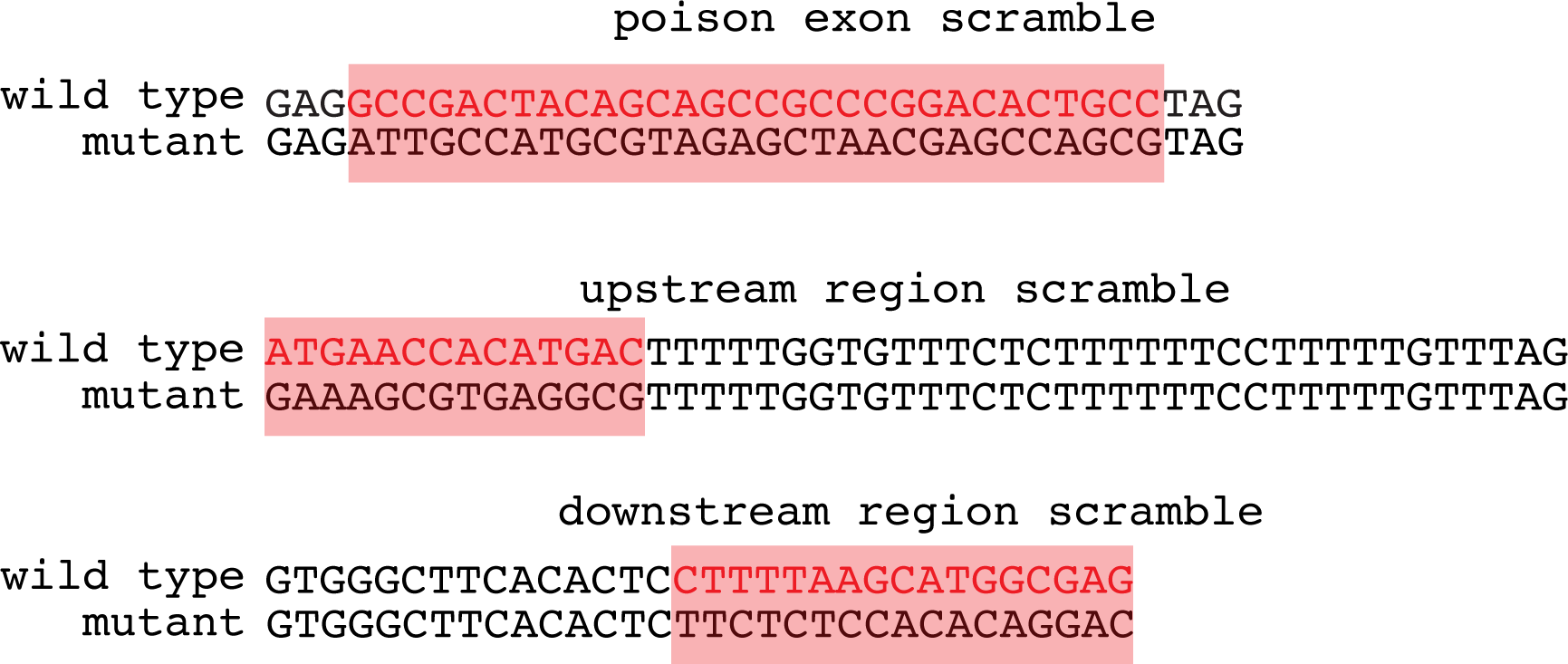
Scramble mutagenesis sequences for. Figure 3D. For the construct used in Figure 3D, wild type and scramble mutagenesis sequences are highlighted.

**Figure S6.**
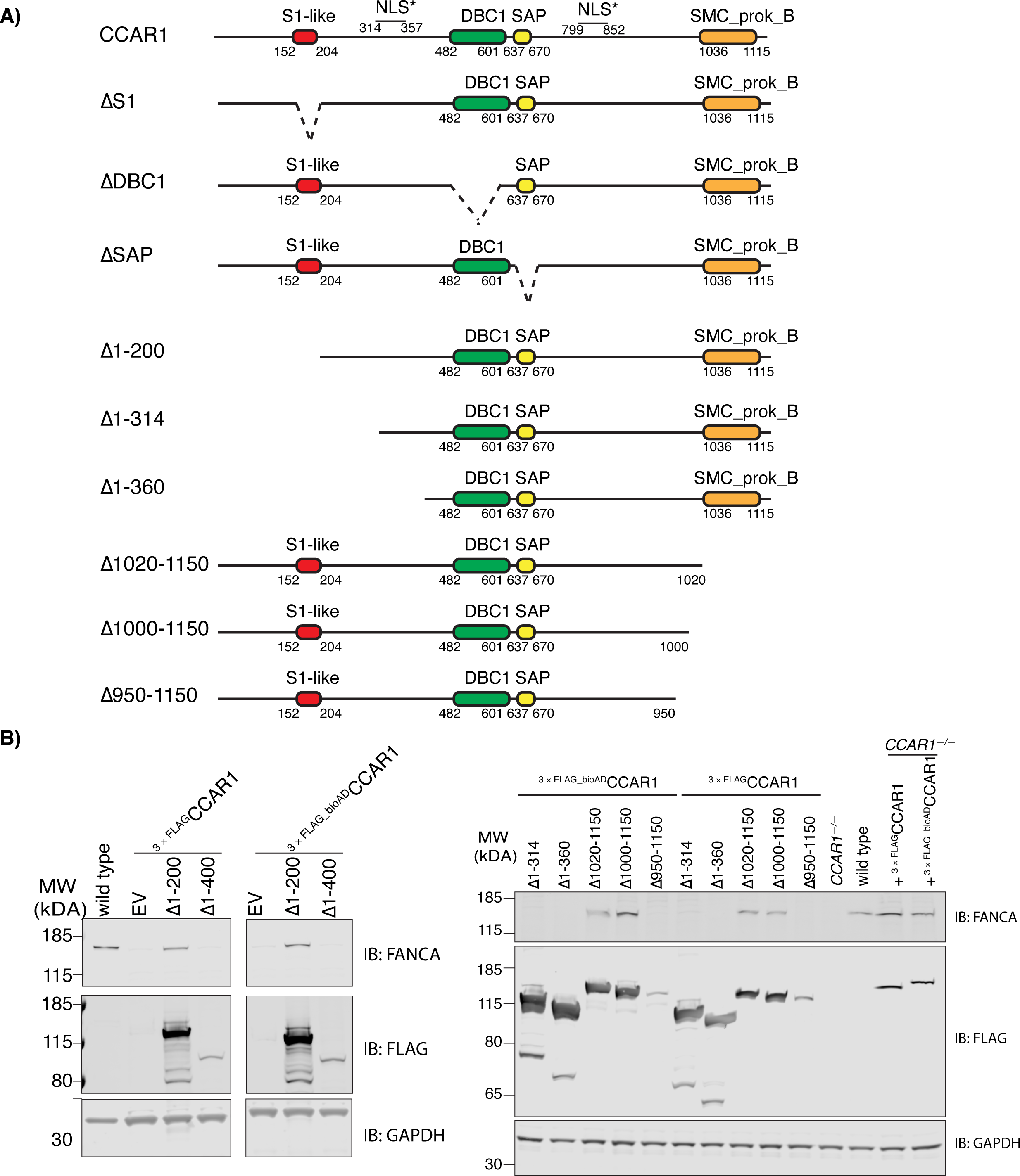
Truncations of N terminal and C terminal CCAR1 construct reveals the minimum functional unit of CCAR1 to suppress FANCA PE exon inclusion. **A)** Schematics of CCAR1 protein constructs tested. **B)** As shown in Figure 3D, *CCAR1^-/-^* K562 cells were transduced with the indicated constructs and analyzed by the Western blotting using antibodies anti-FANCA, anti-FLAG and anti-GAPDH. Deletion of first 400 a.a sequence did not rescue FANCA protein expression, probably due to disruption to predicted NLS sequence. The N terminal deletion was further narrowed down by generating deletion of first 314 a.a (which should not affect the predicted NLS sequences) or 368 a.a. However, even first 314 a.a sequence did not rescue FANCA expression. On the C terminal deletion experiments showed that last 150 a.a sequence was dispensable for the FANCA PE suppression, but deleting last 200 a.a sequence (1-950 a.a) did not rescue the FANCA expression.

**Figure S7.**
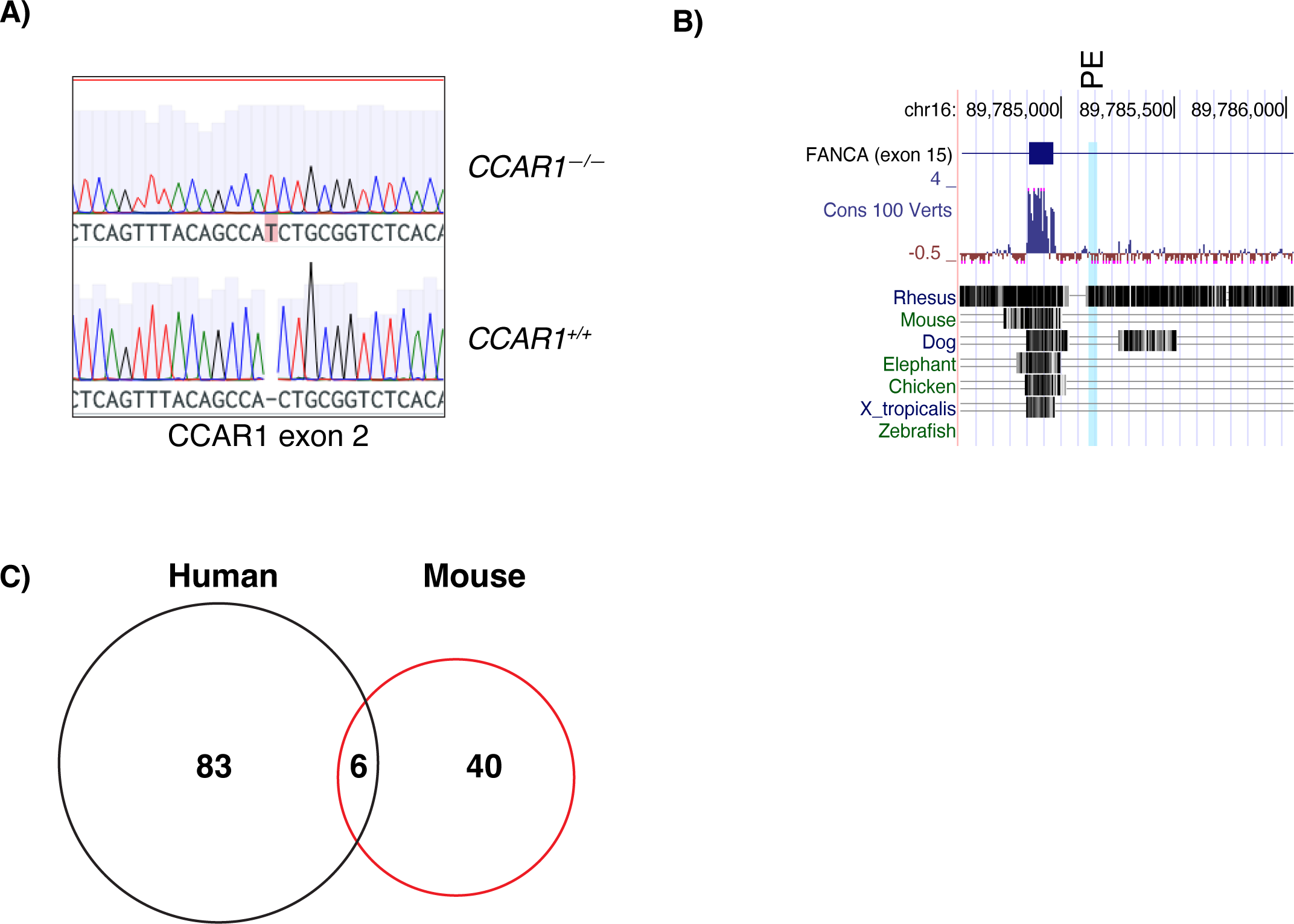
The FANCA poison exon is not evolutionarily conserved. **A)** Sanger sequencing of the mouse *Ccar1* ­3T3 knockout clone. The Sanger tracks indicate a homozygous 1 bp T insertion at the exon 2 of *Ccar1* in mouse 3T3 cells. **B)** The region surrounding exon 15 of *FANCA* was analyzed in USCS genome browser. The conservation track (MultiViz-470) was used to visualize whether the sequences are conserved. Except Rhesus Monkey, other speciess lack the site containing the poison exon. **C)** Venn diagram showing genes with splicing affected by *CCAR1* depletion in human and mouse.

**Figure S8.**
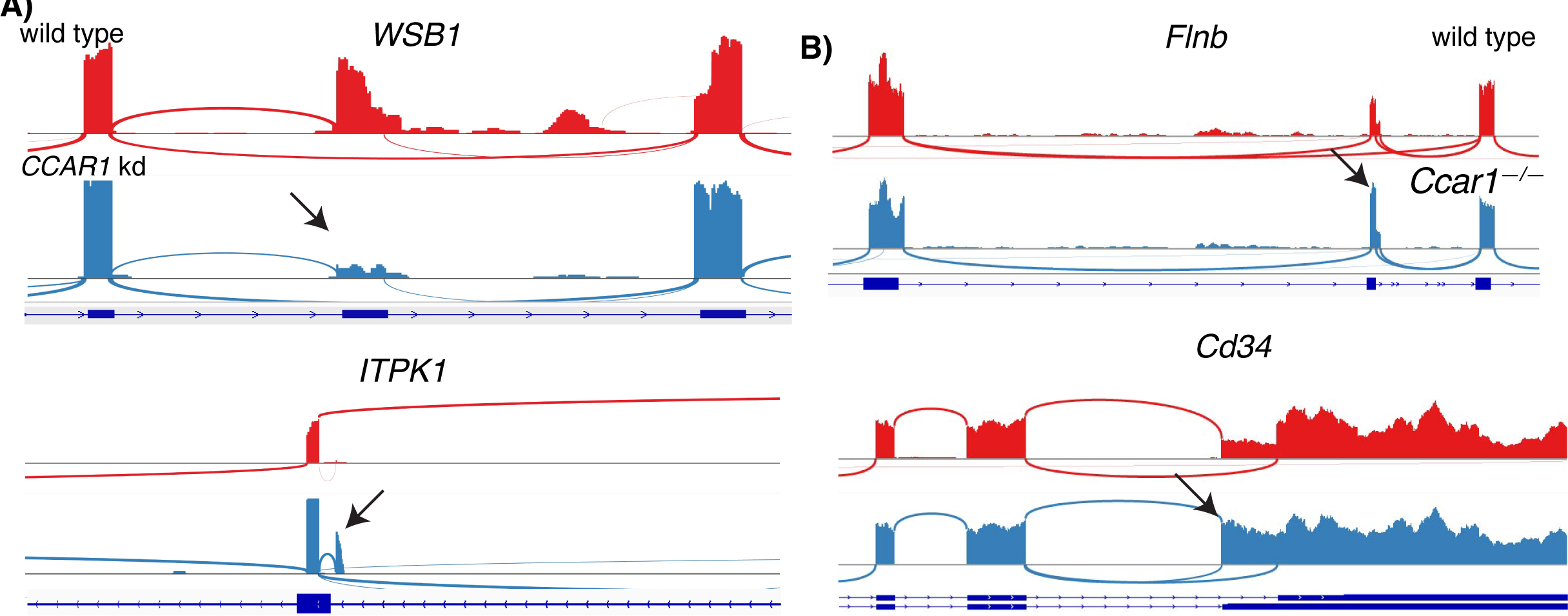
Examples of differential alternative splicing in *CCAR1* depleted cells in human and mouse cell lines. **A)** Sashimi plots show different alternative splicing events in human cells **B)** Sashimi plots show different alternative splicing events in mouse cells

**Figure S9.**
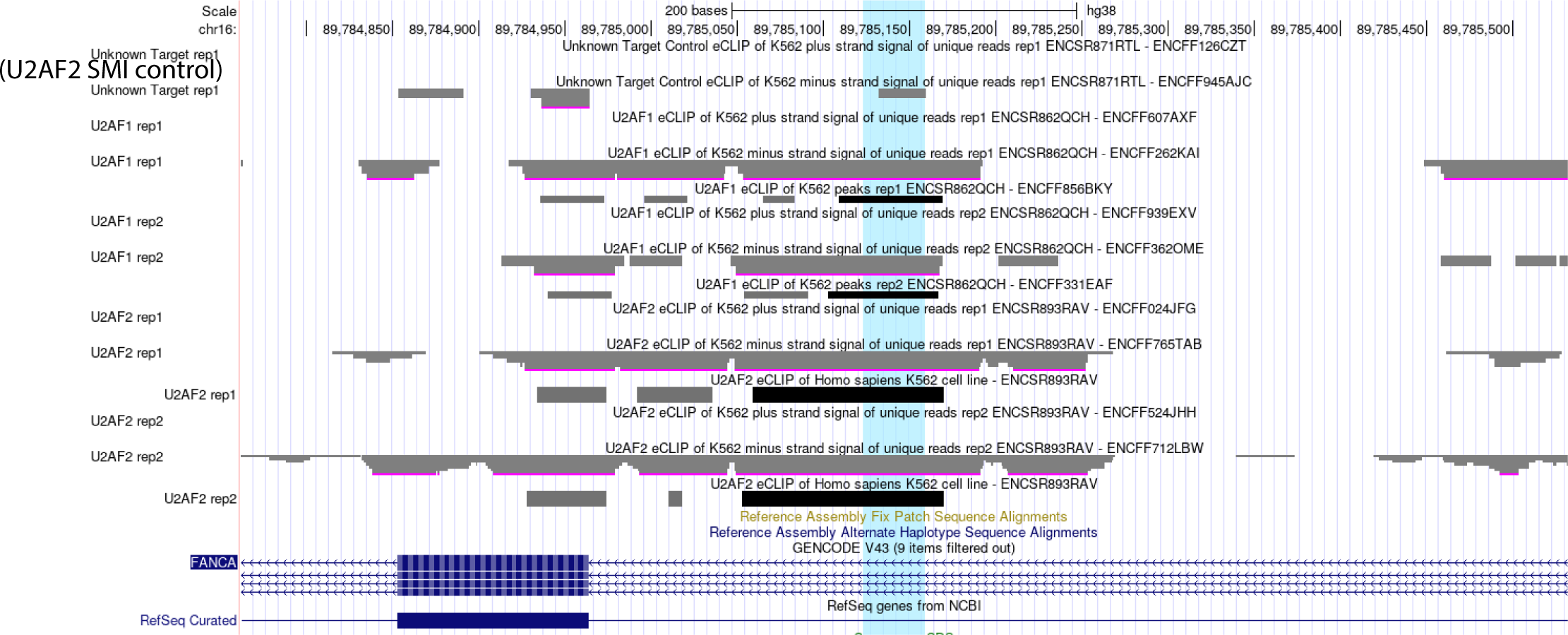
eCLIP-Seq PEAKs from U2AF1 and U2AF2 around the FANCA poison exon inclusion site. eCLIP ENCODE data sets of multiple splicing factors ^44^ are visualized around the *FANCA* poison exon (highlighted light blue). Size matched input from U2AF2 samples was used as a control.

## Reference

1. Molinaro, C., Martoriati, A., and Cailliau, K. (2021). Proteins from the DNA Damage Response: Regulation, Dysfunction, and Anticancer Strategies. Cancers (Basel) 13. 10.3390/cancers13153819.

2. Pierce, A.J., Hu, P., Han, M., Ellis, N., and Jasin, M. (2001). Ku DNA end-binding protein modulates homologous repair of double-strand breaks in mammalian cells. Genes Dev. 15, 3237–3242. 10.1101/gad.946401.

3. Cui, X., and Meek, K. (2007). Linking double-stranded DNA breaks to the recombination activating gene complex directs repair to the nonhomologous end-joining pathway. Proc Natl Acad Sci USA 104, 17046–17051. 10.1073/pnas.0610928104.

4. Carballar, R., Martínez-Láinez, J.M., Samper, B., Bru, S., Bállega, E., Mirallas, O., Ricco, N., Clotet, J., and Jiménez, J. (2020). CDK-mediated Yku80 Phosphorylation Regulates the Balance Between Non-homologous End Joining (NHEJ) and Homologous Directed Recombination (HDR). J. Mol. Biol. 432, 166715. 10.1016/j.jmb.2020.11.014.

5. Yeh, C.D., Richardson, C.D., and Corn, J.E. (2019). Advances in genome editing through control of DNA repair pathways. Nat. Cell Biol. 21, 1468–1478. 10.1038/s41556-019-0425-z.

6. Barman, A., Deb, B., and Chakraborty, S. (2020). A glance at genome editing with CRISPR-Cas9 technology. Curr. Genet. 66, 447–462. 10.1007/s00294-019-01040-3.

7. Gabrielli, B.G., Sarcevic, B., Sinnamon, J., Walker, G., Castellano, M., Wang, X.Q., and Ellem, K.A. (1999). A cyclin D-Cdk4 activity required for G2 phase cell cycle progression is inhibited in ultraviolet radiation-induced G2 phase delay. J. Biol. Chem. 274, 13961–13969. 10.1074/jbc.274.20.13961.

8. Oh, J., and Symington, L.S. (2018). Role of the mre11 complex in preserving genome integrity. Genes (Basel) 9. 10.3390/genes9120589.

9. Yang, D., Scavuzzo, M.A., Chmielowiec, J., Sharp, R., Bajic, A., and Borowiak, M. (2016). Enrichment of G2/M cell cycle phase in human pluripotent stem cells enhances HDR-mediated gene repair with customizable endonucleases. Sci. Rep. 6, 21264. 10.1038/srep21264.

10. Huertas, D., Sendra, R., and Muñoz, P. (2009). Chromatin dynamics coupled to DNA repair. Epigenetics 4, 31–42. 10.4161/epi.4.1.7733.

11. Richardson, C.D., Kazane, K.R., Feng, S.J., Zelin, E., Bray, N.L., Schäfer, A.J., Floor, S.N., and Corn, J.E. (2018). CRISPR-Cas9 genome editing in human cells occurs via the Fanconi anemia pathway. Nat. Genet. 50, 1132–1139. 10.1038/s41588-018-0174-0.

12. Wienert, B., Nguyen, D.N., Guenther, A., Feng, S.J., Locke, M.N., Wyman, S.K., Shin, J., Kazane, K.R., Gregory, G.L., Carter, M.A.M., et al. (2020). Timed inhibition of CDC7 increases CRISPR-Cas9 mediated templated repair. Nat. Commun. 11, 2109. 10.1038/s41467-020-15845-1.

13. Karasu, M.E., Toufektchan, E., Maciejowski, J., and Corn, J.E. (2022). TREX1 restricts CRISPR-Cas9 genome editing in human cells. BioRxiv. 10.1101/2022.12.12.520063.

14. Rishi, A.K., Zhang, L., Boyanapalli, M., Wali, A., Mohammad, R.M., Yu, Y., Fontana, J.A., Hatfield, J.S., Dawson, M.I., Majumdar, A.P.N., et al. (2003). Identification and characterization of a cell cycle and apoptosis regulatory protein-1 as a novel mediator of apoptosis signaling by retinoid CD437. J. Biol. Chem. 278, 33422–33435. 10.1074/jbc.M303173200.

15. Kim, J.H., Yang, C.K., Heo, K., Roeder, R.G., An, W., and Stallcup, M.R. (2008). CCAR1, a key regulator of mediator complex recruitment to nuclear receptor transcription complexes. Mol. Cell 31, 510–519. 10.1016/j.molcel.2008.08.001.

16. Olivieri, M., Cho, T., Álvarez-Quilón, A., Li, K., Schellenberg, M.J., Zimmermann, M., Hustedt, N., Rossi, S.E., Adam, S., Melo, H., et al. (2020). A genetic map of the response to DNA damage in human cells. Cell 182, 481–496.e21. 10.1016/j.cell.2020.05.040.

17. Matsushita, N., Kitao, H., Ishiai, M., Nagashima, N., Hirano, S., Okawa, K., Ohta, T., Yu, D.S., McHugh, P.J., Hickson, I.D., et al. (2005). A FancD2-monoubiquitin fusion reveals hidden functions of Fanconi anemia core complex in DNA repair. Mol. Cell 19, 841–847. 10.1016/j.molcel.2005.08.018.

18. Smogorzewska, A., Matsuoka, S., Vinciguerra, P., McDonald, E.R., Hurov, K.E., Luo, J., Ballif, B.A., Gygi, S.P., Hofmann, K., D’Andrea, A.D., et al. (2007). Identification of the FANCI protein, a monoubiquitinated FANCD2 paralog required for DNA repair. Cell 129, 289–301. 10.1016/j.cell.2007.03.009.

19. Johnson, G.S., Rajendran, P., and Dashwood, R.H. (2020). CCAR1 and CCAR2 as gene chameleons with antagonistic duality: Preclinical, human translational, and mechanistic basis. Cancer Sci. 111, 3416–3425. 10.1111/cas.14579.

20. Muthu, M., Cheriyan, V.T., and Rishi, A.K. (2015). CARP-1/CCAR1: a biphasic regulator of cancer cell growth and apoptosis. Oncotarget 6, 6499–6510. 10.18632/oncotarget.3376.

21. Leung, J.W.C., Wang, Y., Fong, K.W., Huen, M.S.Y., Li, L., and Chen, J. (2012). Fanconi anemia (FA) binding protein FAAP20 stabilizes FA complementation group A (FANCA) and participates in interstrand cross-link repair. Proc Natl Acad Sci USA 109, 4491–4496. 10.1073/pnas.1118720109.

22. Garcia-Higuera, I., Kuang, Y., Denham, J., and D’Andrea, A.D. (2000). The fanconi anemia proteins FANCA and FANCG stabilize each other and promote the nuclear accumulation of the Fanconi anemia complex. Blood 96, 3224–3230. 10.1182/blood.V96.9.3224.

23. ENCODE Project Consortium (2012). An integrated encyclopedia of DNA elements in the human genome. Nature 489, 57–74. 10.1038/nature11247.

24. Brunquell, J., Yuan, J., Erwin, A., Westerheide, S.D., and Xue, B. (2014). DBC1/CCAR2 and CCAR1 Are Largely Disordered Proteins that Have Evolved from One Common Ancestor. Biomed Res. Int. 2014, 418458. 10.1155/2014/418458.

25. Bycroft, M., Hubbard, T.J., Proctor, M., Freund, S.M., and Murzin, A.G. (1997). The solution structure of the S1 RNA binding domain: a member of an ancient nucleic acid-binding fold. Cell 88, 235–242. 10.1016/s0092-8674(00)81844-9.

26. Anantharaman, V., and Aravind, L. (2008). Analysis of DBC1 and its homologs suggests a potential mechanism for regulation of sirtuin domain deacetylases by NAD metabolites. Cell Cycle 7, 1467–1472. 10.4161/cc.7.10.5883.

27. Aravind, L., and Koonin, E.V. (2000). SAP - a putative DNA-binding motif involved in chromosomal organization. Trends Biochem. Sci. 25, 112–114. 10.1016/s0968-0004(99)01537-6.

28. Weighardt, F., Cobianchi, F., Cartegni, L., Chiodi, I., Villa, A., Riva, S., and Biamonti, G. (1999). A novel hnRNP protein (HAP/SAF-B) enters a subset of hnRNP complexes and relocates in nuclear granules in response to heat shock. J. Cell Sci. 112 ( *Pt 10*), 1465–1476. 10.1242/jcs.112.10.1465.

29. Nayler, O., Strätling, W., Bourquin, J.P., Stagljar, I., Lindemann, L., Jasper, H., Hartmann, A.M., Fackelmayer, F.O., Ullrich, A., and Stamm, S. (1998). SAF-B protein couples transcription and pre-mRNA splicing to SAR/MAR elements. Nucleic Acids Res. 26, 3542–3549. 10.1093/nar/26.15.3542.

30. Rishi, A.K., Zhang, L., Yu, Y., Jiang, Y., Nautiyal, J., Wali, A., Fontana, J.A., Levi, E., and Majumdar, A.P.N. (2006). Cell cycle- and apoptosis-regulatory protein-1 is involved in apoptosis signaling by epidermal growth factor receptor. J. Biol. Chem. 281, 13188–13198. 10.1074/jbc.M512279200.

31. Vaquero-Garcia, J., Barrera, A., Gazzara, M.R., González-Vallinas, J., Lahens, N.F., Hogenesch, J.B., Lynch, K.W., and Barash, Y. (2016). A new view of transcriptome complexity and regulation through the lens of local splicing variations. eLife 5, e11752. 10.7554/eLife.11752.

32. Raj, B., Irimia, M., Braunschweig, U., Sterne-Weiler, T., O’Hanlon, D., Lin, Z.-Y., Chen, G.I., Easton, L.E., Ule, J., Gingras, A.-C., et al. (2014). A global regulatory mechanism for activating an exon network required for neurogenesis. Mol. Cell 56, 90–103. 10.1016/j.molcel.2014.08.011.

33. Fu, R., Zhu, Y., Jiang, X., Li, Y., Zhu, M., Dong, M., Huang, Z., Wang, C., Labouesse, M., and Zhang, H. (2018). CCAR-1 affects hemidesmosome biogenesis by regulating unc-52/perlecan alternative splicing in the C. elegans epidermis. J. Cell Sci. 131. 10.1242/jcs.214379.

34. Hegele, A., Kamburov, A., Grossmann, A., Sourlis, C., Wowro, S., Weimann, M., Will, C.L., Pena, V., Lührmann, R., and Stelzl, U. (2012). Dynamic protein-protein interaction wiring of the human spliceosome. Mol. Cell 45, 567–580. 10.1016/j.molcel.2011.12.034.

35. Beusch, I., Rao, B., Studer, M.K., Luhovska, T., Šukytė, V., Lei, S., Oses-Prieto, J., SeGraves, E., Burlingame, A., Jonas, S., et al. (2023). Targeted high-throughput mutagenesis of the human spliceosome reveals its in vivo operating principles. Mol. Cell. 10.1016/j.molcel.2023.06.003.

36. Chen, S., Francioli, L.C., Goodrich, J.K., Collins, R.L., Kanai, M., Wang, Q., Alföldi, J., Watts, N.A., Vittal, C., Gauthier, L.D., et al. (2022). A genome-wide mutational constraint map quantified from variation in 76,156 human genomes. BioRxiv. 10.1101/2022.03.20.485034.

37. Lareau, L.F., Inada, M., Green, R.E., Wengrod, J.C., and Brenner, S.E. (2007). Unproductive splicing of SR genes associated with highly conserved and ultraconserved DNA elements. Nature 446, 926–929. 10.1038/nature05676.

38. Leclair, N.K., Brugiolo, M., Urbanski, L., Lawson, S.C., Thakar, K., Yurieva, M., George, J., Hinson, J.T., Cheng, A., Graveley, B.R., et al. (2020). Poison Exon Splicing Regulates a Coordinated Network of SR Protein Expression during Differentiation and Tumorigenesis. Mol. Cell 80, 648–665.e9. 10.1016/j.molcel.2020.10.019.

39. Carvill, G.L., and Mefford, H.C. (2020). Poison exons in neurodevelopment and disease. Curr. Opin. Genet. Dev. 65, 98–102. 10.1016/j.gde.2020.05.030.

40. Rogalska, M.E., Vivori, C., and Valcárcel, J. (2023). Regulation of pre-mRNA splicing: roles in physiology and disease, and therapeutic prospects. Nat. Rev. Genet. 24, 251–269. 10.1038/s41576-022-00556-8.

41. Yu, G. (2020). Gene ontology semantic similarity analysis using gosemsim. Methods Mol. Biol. 2117, 207–215. 10.1007/978-1-0716-0301-7_11.

42. Wu, T., Hu, E., Xu, S., Chen, M., Guo, P., Dai, Z., Feng, T., Zhou, L., Tang, W., Zhan, L., et al. (2021). clusterProfiler 4.0: A universal enrichment tool for interpreting omics data. Innovation (Camb) 2, 100141. 10.1016/j.xinn.2021.100141.

43. Van Nostrand, E.L., Freese, P., Pratt, G.A., Wang, X., Wei, X., Xiao, R., Blue, S.M., Chen, J.-Y., Cody, N.A.L., Dominguez, D., et al. (2020). A large-scale binding and functional map of human RNA-binding proteins. Nature 583, 711–719. 10.1038/s41586-020-2077-3.

44. Van Nostrand, E.L., Pratt, G.A., Shishkin, A.A., Gelboin-Burkhart, C., Fang, M.Y., Sundararaman, B., Blue, S.M., Nguyen, T.B., Surka, C., Elkins, K., et al. (2016). Robust transcriptome-wide discovery of RNA-binding protein binding sites with enhanced CLIP (eCLIP). Nat. Methods 13, 508–514. 10.1038/nmeth.3810.

45. Dewitt, M., Wong, J., Wienert, B., Schlapansky, M.F., and Aird, E. (2023). In vitro transcription of guide RNAs and 5’-triphosphate removal.

46. Gibson, D.G., Young, L., Chuang, R.-Y., Venter, J.C., Hutchison, C.A., and Smith, H.O. (2009). Enzymatic assembly of DNA molecules up to several hundred kilobases. Nat. Methods 6, 343–345. 10.1038/nmeth.1318.

47. Parrott, M.B., and Barry, M.A. (2000). Metabolic biotinylation of recombinant proteins in mammalian cells and in mice. Mol. Ther. 1, 96–104. 10.1006/mthe.1999.0011.

48. Gilbert, L.A., Horlbeck, M.A., Adamson, B., Villalta, J.E., Chen, Y., Whitehead, E.H., Guimaraes, C., Panning, B., Ploegh, H.L., Bassik, M.C., et al. (2014). Genome-scale CRISPR-mediated control of gene repression and activation. Cell 159, 647–661. 10.1016/j.cell.2014.09.029.

49. Li, W., Xu, H., Xiao, T., Cong, L., Love, M.I., Zhang, F., Irizarry, R.A., Liu, J.S., Brown, M., and Liu, X.S. (2014). MAGeCK enables robust identification of essential genes from genome-scale CRISPR/Cas9 knockout screens. Genome Biol. 15, 554. 10.1186/s13059-014-0554-4.

50. Colic, M., Wang, G., Zimmermann, M., Mascall, K., McLaughlin, M., Bertolet, L., Lenoir, W.F., Moffat, J., Angers, S., Durocher, D., et al. (2019). Identifying chemogenetic interactions from CRISPR screens with drugZ. Genome Med. 11, 52. 10.1186/s13073-019-0665-3.

51. Kim, D., Langmead, B., and Salzberg, S.L. (2015). HISAT: a fast spliced aligner with low memory requirements. Nat. Methods 12, 357–360. 10.1038/nmeth.3317.

52. Liao, Y., Smyth, G.K., and Shi, W. (2013). The Subread aligner: fast, accurate and scalable read mapping by seed-and-vote. Nucleic Acids Res. 41, e108. 10.1093/nar/gkt214.

53. Love, M.I., Huber, W., and Anders, S. (2014). Moderated estimation of fold change and dispersion for RNA-seq data with DESeq2. Genome Biol. 15, 550. 10.1186/s13059-014-0550-8.

54. Robinson, J.T., Thorvaldsdóttir, H., Winckler, W., Guttman, M., Lander, E.S., Getz, G., and Mesirov, J.P. (2011). Integrative genomics viewer. Nat. Biotechnol. 29, 24–26. 10.1038/nbt.1754.

55. Mironov, A., Petrova, M., Margasyuk, S., Vlasenok, M., Mironov, A.A., Skvortsov, D., and Pervouchine, D.D. (2023). Tissue-specific regulation of gene expression via unproductive splicing. Nucleic Acids Res. 51, 3055–3066. 10.1093/nar/gkad161.

56. Dobin, A., Davis, C.A., Schlesinger, F., Drenkow, J., Zaleski, C., Jha, S., Batut, P., Chaisson, M., and Gingeras, T.R. (2013). STAR: ultrafast universal RNA-seq aligner. Bioinformatics 29, 15–21. 10.1093/bioinformatics/bts635.

57. Frankish, A., Diekhans, M., Jungreis, I., Lagarde, J., Loveland, J.E., Mudge, J.M., Sisu, C., Wright, J.C., Armstrong, J., Barnes, I., et al. (2021). GENCODE 2021. Nucleic Acids Res. 49, D916–D923. 10.1093/nar/gkaa1087.

58. Graubert, A., Aguet, F., Ravi, A., Ardlie, K.G., and Getz, G. (2021). RNA-SeQC 2: efficient RNA-seq quality control and quantification for large cohorts. Bioinformatics 37, 3048–3050. 10.1093/bioinformatics/btab135.

